# Ageing impacts extracellular matrix turnover and remodelling in the kidney

**DOI:** 10.64898/2026.03.02.709057

**Authors:** Rebecca Preston, Anna Hoyle, Alana Stevenson Harris, Emily Williams, Tess Birtles, Joan Chang, Joe Swift, Alexander Eckersley, Rachel Lennon

## Abstract

At least 10% of the global population is impacted by chronic kidney disease (CKD) and ageing is a key risk factor. CKD is characterised by the build-up of extracellular matrix and a loss of functional nephrons. However, the mechanisms that maintain matrix homeostasis across the physiological lifespan remain elusive. Using ¹³C-lysine metabolic labelling, we quantified kidney matrix protein turnover in healthy mice at four timepoints (8, 22, 52, and 78 weeks). We found that basement membrane components, including collagen IV, laminin-521, nidogens and perlecan, were more long-lived over age, with collagen IV half-lives extending from weeks in young kidneys to years in aged kidneys, suggesting a reduced capacity for basement membrane renewal. The half-lives of fibrillar collagens I and III also increased over age up to forty-fold, which is consistent with minimal degradation. In contrast, collagen XV retained rapid turnover despite increased abundance, indicating a persistent role in tissue remodelling. Using peptide location fingerprinting to predict structural alterations and proteolytic processing we identified age-dependent meprin oligomerisation and altered nidogen–laminin interaction states. We predicted structural alterations within assembly domains of collagen VI and reduced accessibility of integrin-binding regions, suggesting altered microfibril organisation and cell-surface binding. Collagen XV had predicted structural changes across the NC1 domain encoding the matrikine restin, consistent with altered protease accessibility and matrikine release during ageing. These findings indicate that age-related kidney fibrosis is primarily caused by impaired matrix degradation, with protease accessibility and altered matrix interactions likely playing key roles in this remodeling process.

## Introduction

Extracellular matrix in the kidney is dynamic and undergoes remodelling throughout life to enable growth and maintain normal tissue architecture and integrity [1]. Beyond providing structural support, the kidney matrix acts as a reservoir for growth factors and proteases [1, 2]. Matrix synthesis and turnover requires tight regulation for maintenance of organ function and matrix accumulation, as occurs in fibrosis, is the common pathological feature of chronic kidney disease (CKD), irrespective of aetiology [3]. CKD represents a major public health concern, affecting over 10% of the global population [4, 5]. Advanced age is a major risk factor for CKD progression and is independently associated with glomerulosclerosis, tubular atrophy and interstitial fibrosis [6–8]. With every country in the world experiencing increased life-expectancy, the risk of developing CKD is projected to double by 2050 [9]. As there are no current effective therapies to prevent or reverse kidney fibrosis, there is an urgent need to understand the mechanisms driving kidney matrix accumulation with age.

To understand matrix dysregulation in ageing and disease, it is a prerequisite to understand matrix homeostasis in healthy tissues. Mass spectrometry-based proteomic studies have characterised matrix composition in many tissues [10–12], and MatrisomeDB, is a comprehensive community resource with the most complete collection of matrix proteomic data [13]. Here, the matrisome is classified in two groups: the core matrisome (collagens, glycoproteins, and proteoglycans); and the associated matrisome (matrix regulators, affiliated proteins, and secreted factors). Another community resource for matrix research is, ‘Basement membraneBase’, which provides a catalogue of proteins that localise or are predicted to localise to basement membranes and adjacent cell surfaces [14].

In the kidney, matrix components organise into basement membranes and looser interstitial matrices which have different functional roles. The glomerular basement membrane (GBM) separates endothelial cells from overlying podocytes and selectively controls glomerular filtration. Major components of the mature GBM include type IV collagen (primarily α3-α4-α5 heterotrimers), laminins (mainly α5β2γ1), nidogens and the heparin sulfate proteoglycans (HSPGs), agrin and perlecan [15]. The mesangial matrix provides structural support for glomerular capillaries and comprises fibronectin, collagens type IV and V, laminins and HSPGs [16]. The Bowman’s capsule surrounds the glomerular capillaries and basement membranes of the capsule are rich in collagen IV α1-α1-α2 and α5-α5-α6 heterotrimers [16, 17]. The looser tubulointerstitial matrix consists of additional collagens (types I, III, IV, V, VI, VII and VIII), glycosaminoglycans, and glycoproteins including fibronectin, versican, biglycan and decorin [16]. Decorin and biglycan are small leucine-rich proteoglycans that are either matrix-bound or released as soluble mediators [18–21]. In healthy kidneys, they are expressed in the interstitium [22] but show de novo glomerular expression in animal models of fibrosis and in human kidney disease [23, 24]. Collagens XV and XVIII belong to the multiplexin collagen superfamily and associate with fibrillar collages near basement membrane zones [25, 26]. These hybrid collagen-proteoglycans contain chondroitin sulfate or heparin sulfate chains and harbour C-terminal domains, restin and endostatin, which upon cleavage exert anti-angiogenic effects [27].

The kidney matrix undergoes balanced remodelling in response to tissue growth and injury. In ageing and disease, this balance is disrupted, leading to excessive matrix production without reciprocal degradation, and in turn, aberrant accumulation [28]. Mechanisms behind abnormal matrix remodelling involve dysregulation of proteases and their inhibitors, including matrix metalloproteinases, ADAMTS proteases, meprins, serine proteases and cathepsins [29]. Proteolytic fragmentation of matrix proteins releases bioactive fragments, known as matrikines, which modulate multiple cellular responses implicated in the pathogenesis of age-related kidney diseases (reviewed in [30]). Many kidney basement membrane components are vulnerable to proteolytic fragmentation, and aberrant matrix remodelling in kidney ageing likely releases diverse matrikines with yet, undetermined roles [30].

Mass spectrometry combined with fractionation techniques, allowing for matrix-enrichment [31], defined kidney matrix composition in health [2, 32–34] and disease [14, 35–37] and comparison of mouse models of kidney disease and human ageing, revealed a common signature of altered matrix [38]. This study demonstrated an overall reduction in basement membrane components and an increase in interstitial matrix proteins. Further integration of 3D imaging with proteomics identified compositional changes in the interstitial matrix and basement membranes during kidney development [39]. Collectively, these studies demonstrate the dynamics of kidney matrix during development and disease and suggest that switches between basement membrane and interstitial matrix proteins could contribute to age associated dysfunction [40]. However, studies focussed on kidney matrix dynamics across the physiological lifespan have not been performed, and mechanisms driving altered matrix composition remain unclear.

Whilst traditional proteomic studies provide snapshots of protein abundance, they do not capture the dynamic processes of protein turnover within tissues. Protein turnover, the balance between protein synthesis and degradation (proteostasis), is fundamental to tissue maintenance [41, 42] and perturbed proteostasis is a hallmark of ageing [43, 44]. During ageing, matrix components become damaged through post-translational modification and cross-linking, leading to matrix accumulation and increased stiffness [45]. Basement membranes have long been considered stable and rigid scaffolds, with components having long half-lives in the order of weeks [46, 47]. However studies in invertebrates (*Drosophila* and *C. elegans*) have demonstrated a far more dynamic matrix with live imaging of fluorescently tagged basement membrane components demonstrating rapid turnover with half-lives as short as 7–10 hours [48, 49]. Whilst imaging of fluorophore-tagged matrix in vertebrate systems is emerging as a tool to measure turnover [50], metabolic labelling combined with mass spectrometry offers powerful means to measure global protein turnover in mammalian systems. Introduction of stable-isotope diets, such as ^15^N or ^13^C lysine, allow quantification of newly synthesised versus pre-existing proteins. Whilst early ^15^N-based studies estimated global turnover in kidney tissue [51], the approach was limited by broad isotope distributions and metabolic recycling [52], hindering the precision of half-life calculations. Conversely, ^13^C-lysine labelling provides higher specificity, allowing precise turnover measurements of many more individual proteins based on predictable mass shifts in proteolytic peptides. This strategy has been successfully applied in mammalian tissues including liver, brain, cartilage, and skin [53–57], however proteome-wide turnover data across the kidney lifespan is not yet available.

Another approach for predicting matrix remodelling events is peptide location fingerprinting (PLF), a proteomic analysis tool capable of detecting structure-associated alterations within protein sequences by analysing regional changes in peptide coverage. Rather than quantifying whole-protein abundance, PLF identifies localised differences in peptide yield (trypsin accessibility) which, when correlated to regions of interaction, cleavage sites and functional domains, is often indicative of proteolytic cleavage, confirmational change, or post-translational damage modifications [58]. This technique has proven particularly valuable in matrix biology, where structural matrix proteins in diverse tissues are frequently subjected to selective enzymatic processing [59–62]. In the context of kidney health, PLF was applied to investigate matrix remodelling in aged versus young human kidney [63], identifying structure-associated alterations within basement membrane components predicting disrupted molecular interactions, proteolytic degradation, oxidative damage, and altered matrikine release. Determining which kidney matrix proteins are susceptible to structure-associated changes and elucidating upstream mechanisms and downstream consequences, is key to understand age-related kidney degeneration and identify candidates for therapeutic intervention.

Applying these tools to study kidney ageing, we performed a deep proteomic analysis of kidney matrix turnover through development and ageing. Using ¹³C-lysine labelling we aimed to generate proteome-wide turnover data to characterise how matrix remodelling changes across the lifespan and understand how ageing influences basement membrane stability and proteolytic activity. Our findings demonstrate coordinated, yet divergent, matrix dynamics in development and ageing, revealing that disrupted proteolytic degradation rather than excessive synthesis underlies age-related kidney fibrosis.

## Results

### Heavy isotope pulse-labelling of the mouse kidney matrisome

To assess protein turnover in the adult kidney across the lifespan, we conducted heavy-isotope pulse labelling by administering a ^13^C-Lys (heavy diet) for four weeks prior to sacrifice (**Figure 1A**). We analysed kidney tissue from mice sacrificed at four different ages: 8-weeks (young), 22-weeks (adult), 52-weeks (old) and 78-weeks (aged). Kidney tissue was fractionated using a two-step protocol (see **Methods**), yielding a soluble cellular fraction (F1) and a matrix-enriched fraction (F2) (**Figure 1B**). We calculated heavy-to-light ratios for each age group from the heavy and light protein abundances, reflecting ^13^C-Lys incorporation into kidney proteins. Increased ^13^C-Lys incorporation results in increased heavy-to-light ratios, translating to increased turnover rates and shorter half-lives. Combined heavy-to-light ratios from F1 and F2 were highest in 8-week mice and decreased with maturation to 22-weeks, confirming increased protein turnover during development. Protein turnover then increased with advancing age. Within the kidney matrisome (F2 only), this pattern was generally preserved, however matrix protein turnover began to decline in the oldest group (78 weeks). To focus our analysis on the effects of development and ageing on matrix protein abundance and turnover, we performed a comparative analysis using the matrix-enriched (F2) fraction across the lifespan. Principal components analysis (PCA) of F2 revealed age-dependent segregation of samples (**Supplementary Figure 1A&B**). We identified 124, 137, 142 and 153 matrix proteins in F2 for the 8-week, 22-week, 52-week and 78-week mice, respectively and matrix enrichment in F2 increased with advancing age, indicating decreased kidney matrix solubility (**Figure 1D**).

**Figure 1:**
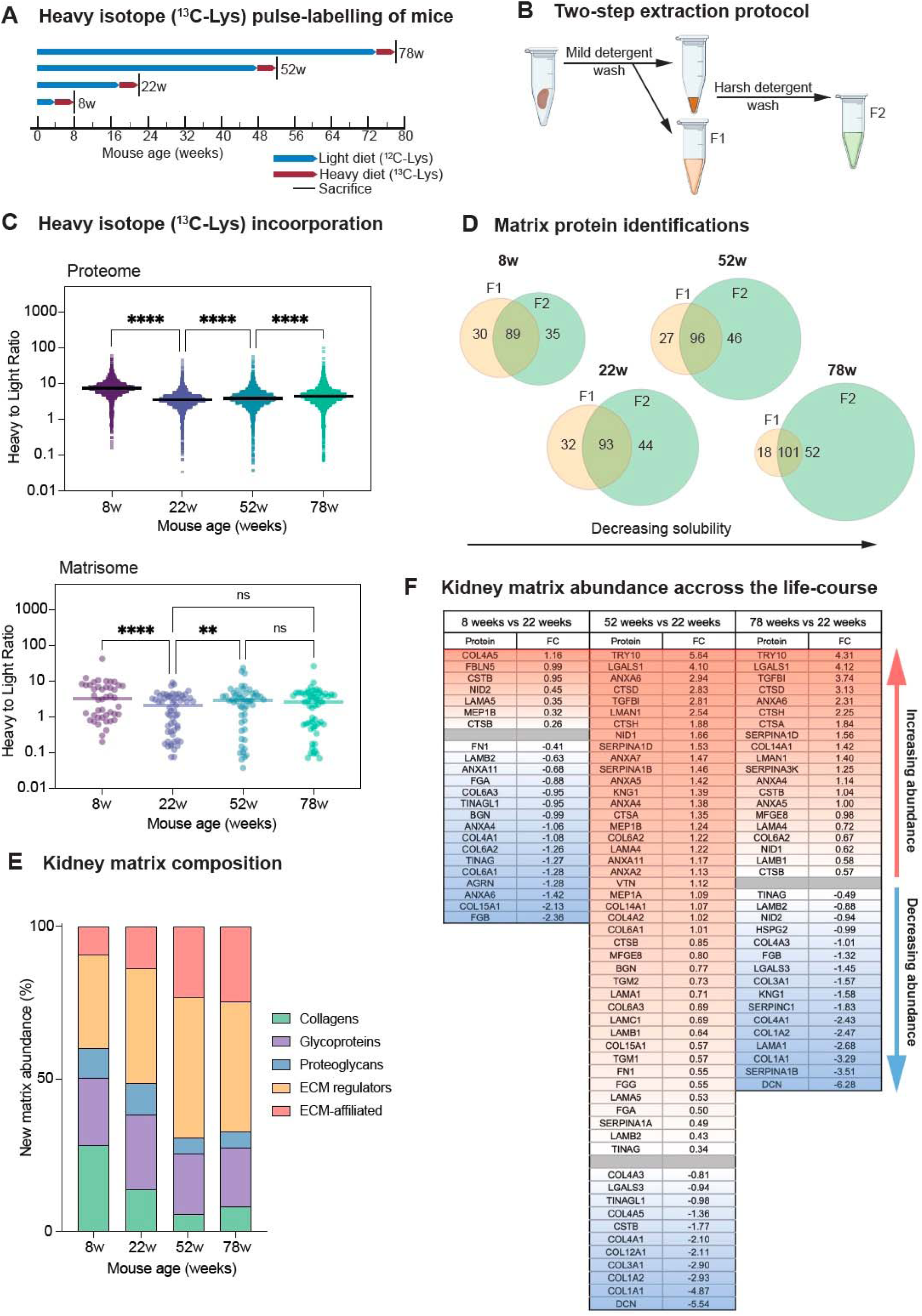
Heavy isotope pulse-labelling and compositional shifts of the mouse kidney matrisome. **A**) Experimental design for heavy isotope (^13^C-Lys) pulse-labelling used for half-life estimation. Mice were fed 4-week pulses of heavy diet (red bars) until they were sacrificed at 8-weeks, 22-weeks, 52-weeks, or 78-weeks (black bars). **B)** Schematic demonstrating the two-step extraction protocol used for fractionation and matrix-enrichment of mouse kidney. **C)** Upper panel: heavy-to-light ratios for all proteins (F1 & F2) detected in kidney tissue from mice fed a ^13^C-Lys diet for 4-weeks at different ages showing decreased ^13^C-Lys incorporation at 22-weeks compared to 8-weeks and then increased ^13^C-Lys incorporation with increasing age. ****p=<0.0001 by Kruskal-Wallis test with correction for multiple comparisons. Bars show median. Lower panel: heavy-to-light ratios for matrix proteins (F2 only) detected in kidney tissue from mice fed ^13^C-Lys diet for 4-weeks showing decreased ^13^C-Lys incorporation from 8-weeks to 22-weeks and increased ^13^C-Lys incorporation from 22-weeks to 52-weeks, which was not sustained with advanced age at 78-weeks. ****p=<0.0001, **p=0.0013 by Kruskal-Wallis test with correction for multiple comparisons. Bars show median. **D**) Venn diagrams illustrating matrix protein identifications in each fraction at 8-, 22-, 52- and 78-weeks, demonstrating increased matrix protein enrichment and decreased matrix solubility with advancing age. **E**) Distribution of new kidney matrix abundance derived from heavy MS1 ion intensities from matrisome proteins in the matrix-enriched fraction (F2) from 8-, 22-, 52- and 78-week-old mice demonstrating a relative reduction in core matrix proteins with age. **F**) Table showing significantly differentially expressed matrix proteins identified in the matrix-enriched fraction from 8-week (left), 52-week (middle) and 78-week mice (right), compared to 22-weeks. Red is upregulated and blue is downregulated based on FC (fold change) in normalised abundance between each condition. Significance threshold determined using adjusted p-value <0.05. N=3 biological replicates per condition for all analysis.

### Kidney matrix composition is altered across the lifespan

To identify compositional changes in the kidney matrix across the lifespan, we calculated total abundance of matrix proteins by combining heavy and light MS1 ion intensities and to estimate new matrix production we used the heavy MS1 ion intensities alone (Proteome Discoverer SILAC workflow). We then analysed protein distribution based on MatrisomeDB categories [13]. Overall, core matrix protein abundance decreased slightly with age, particularly collagens which decreased from 38% and 39% in 8-22 week mice to 27-29% in 52-78 week mice (**Supplementary Figure 1C**). This corroborates previous studies showing increased matrix deposition during development [39]. Interestingly, new matrix production showed substantial changes. 28% of new matrix in 8-week mice was collagens, decreasing to 13% at 22 weeks and further to 5-8% in 52-78 weeks (**Figure 1E**). Significant differential expression was revealed during development (8-week vs. 22-week) and ageing (52-week and 78-week vs. 22-week (**Figure 1F and Supplementary Figures 1D-E**). Consistent with previous findings [38], we observed a compositional shift towards an increased fibrillar matrix with age. There was decreased abundance of collagen XV in 8-week mice but significant increases by 52 weeks. Conversely, decorin (DCN), a proteoglycan that inhibits TGF-β and matrix accumulation [64], was the most notably decreased matrix protein in 52-week and 78-week mice. Both proteins are linked to kidney fibrosis and represent potential therapeutic targets [16, 24, 65]. We also noted shifts in basement membrane composition and kidney protease activity. During development, basement membrane components (COL4A5, NID2, LAMA5) kidney proteases (CTSB, MEP1A) and the protease inhibitor CSTB were more abundant, indicating active basement membrane remodelling during matrix deposition. In aged kidneys (78 weeks), collagen abundance generally decreased with the exception of collagens VI and XIV. Similarly, basement membrane components (collagen IV, LAMB2, NID2, HSPG2, TINAG) decreased, whilst NID1, LAMB1, and LAMA increased. This pattern aligns with previous analyses of aged human kidney [38], suggesting conserved compositional shifts and loss of equilibrium between matrix synthesis and degradation with ageing.

### Age-dependent changes in kidney matrix turnover across the lifespan

To further examine the underlying mechanisms of kidney matrix accumulation we next quantified matrix protein turnover. Heavy-to-light ratios were converted to half-lives for hundreds of kidney matrix proteins, using an established mathematical modelling pipeline (see **Methods**) [66]. To assess alterations in turnover rates across the lifespan, we calculated fold-change (FC) in half-lives (with adjusted p<0.05) during development (8-week vs. 22-week) and ageing (52-week and 78-week vs. 22-week) (**Figure 2A**). In young mice (8 weeks), kidney matrix proteins displayed significantly shorter half-lives compared to adult mice (22 weeks), indicating accelerated turnover during post-natal kidney maturation (**Supplementary Figure 2A**). This rapid turnover encompassed both basement membrane components and interstitial collagens, reflecting active matrix remodelling during late kidney development. By 22 weeks, matrix protein turnover had slowed substantially, indicating a more stable matrix. In older kidneys turnover slowed further, with most core matrix proteins demonstrating significantly increased half-lives at 52 and 78 weeks, compared to 22 weeks (**Figure 2A; Supplementary Figure 2A-C**). This global reduction spanned multiple matrix groups, including basement membrane components and interstitial matrix. Notably, collagen XV displayed distinct expression and turnover dynamics, representing one of few matrix proteins with significantly increased turnover with age, reflected by a decrease in estimated half-life from 153 days in 22-week mice to just 7-9 days in older mice (52 and 78 weeks). This indicates that collagen XV remains dynamically remodelled in the ageing kidney, potentially reflecting tightly regulated synthesis and degradation during age-associated matrix reorganisation. Basement membrane components, including collagen IV α-chains, laminin-521 chains, perlecan, and nidogens, demonstrated sustained decreases in turnover from development and throughout ageing (**Figure 2B**). Collagen IV half-lives increased from weeks in young mice (median 21 days at 8 weeks) to months in adult mice (median 59 days at 22 weeks), before extending to years in aged mice (52 and 78 weeks). Interestingly, collagen IV-α2 displayed notably shorter half-lives during ageing compared to other collagen IV chains. Basement membrane laminin chains similarly showed pronounced half-life increases with age. Laminin- α5, laminin-β2, and laminin-γ1, which form the laminin-521 heterotrimer, all exhibited significantly reduced turnover in aged mice (52 and 78 weeks), with half-lives increasing from 1-2 months at 22 weeks to 3-5 months by 52 and 78 weeks. This indicates that substantial stabilisation of the basement membrane occurs by early adulthood and plateaus during ageing, potentially limiting later capacity for basement membrane renewal or repair. Nidogens and perlecan showed similar age-related reduction in turnover but with more modest changes, suggesting differential regulation of basement membrane proteoglycans compared to collagen IV and laminin networks.

**Figure 2:**
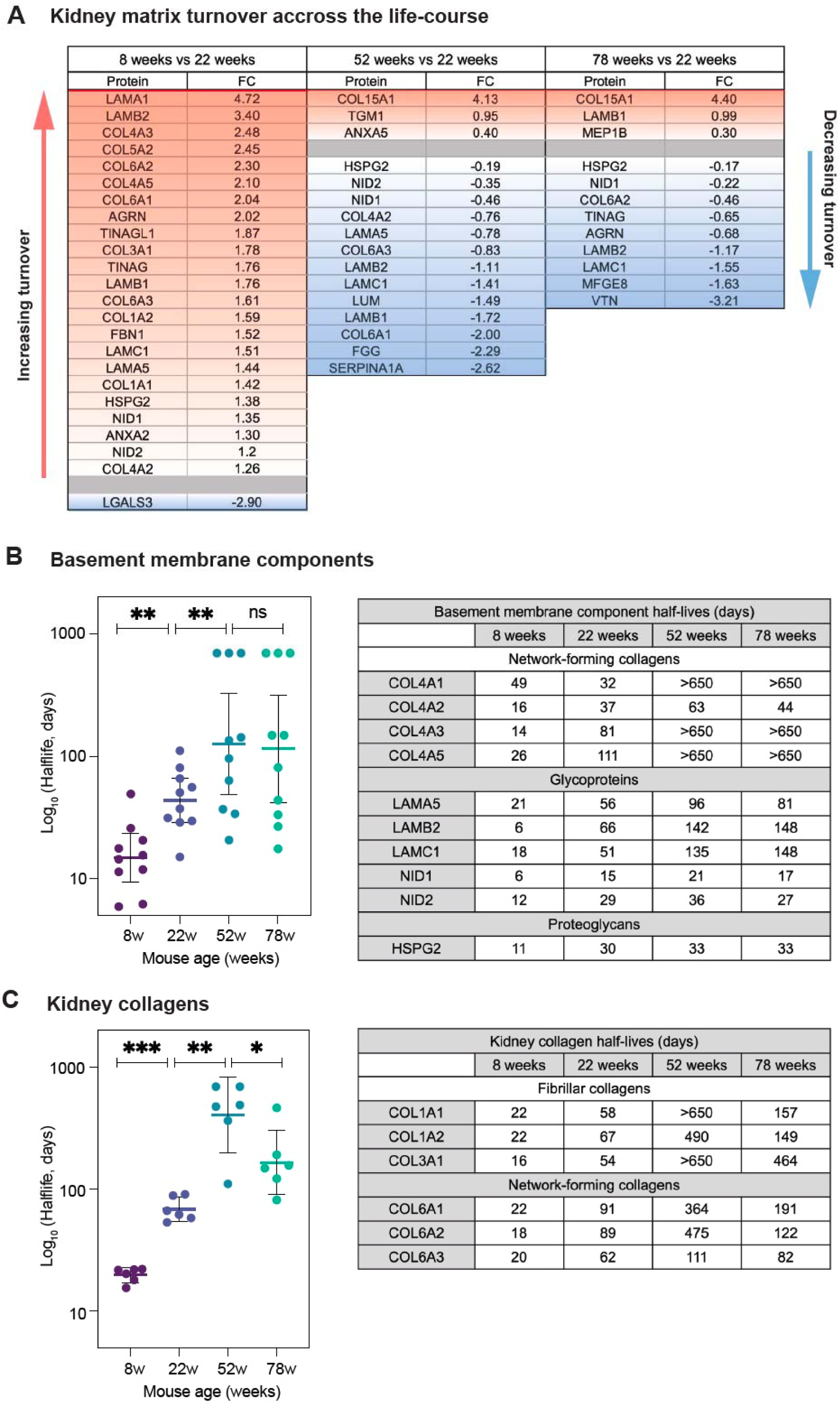
Age-dependent changes in kidney matrix turnover across the lifespan. **A**) Table showing significantly differentially turned over matrix proteins identified in the matrix-enriched fraction from 8-week (left), 52-week (middle) and 78-week mice (right), compared to 22-weeks. Red is upregulated and blue is downregulated based on FC (fold change) in protein half-lives between each condition. Significance threshold determined using adjusted p-value <0.05. **B**) Left panel: half-life estimates for BM components detected in the matrix-enriched fraction from 8-week mice were significantly shorter than 22-week mice whilst half-life estimates were significantly longer in 52-week mice. **p=0.0098, **p=0.0020 by Wilcoxon matched pairs signed rank test. Bars show median. Right panel: half-life estimates (days) of specific BM components, including Col4, laminins, nidogens and perlecan in kidney from across the lifespan. **C**) Left panel: half-life estimates for kidney collagens detected in the matrix-enriched fraction from 8-week mice were significantly shorter than 22-week mice whilst half-life estimates were significantly longer in 52-week and 78-week mice. ***p=0.0005, **p=0.0072, *p=0.0115 by Wilcoxon matched pairs signed rank test. Bars show median. Right panel: half-life estimates (days) of kidney collagens, including fibrillar and network-forming collagens in kidney from across the lifespan. N=3 biological replicates per condition for all analysis.

Interstitial collagens exhibited distinct turnover dynamics across the lifespan (**Figure 2C**). Fibrillar collagens (COL1A1, COL1A2, COL3A1) had rapid turnover in young mice with half-lives of 10-22 days. At 22 weeks, these half-lives increased to 54-67 days, before further extending to over 400 days by 52 weeks. Beaded filament collagen chains (COL6A1, COL6A2, COL6A3) showed similar patterns although were notably less long-lived at 52 weeks. By 78 weeks, these kidney collagens demonstrated reversal in rate of turnover, with half-lives reducing compared to 52 weeks. This suggests potential reversal of collagen cross-linking in advanced age, allowing for increased collagen degradation. Interestingly, the kidney collagen cross-linking enzyme tissue transglutaminase 2 (TGM2) [67] was significantly increased in abundance at 52 weeks when collagen half-lives peaked but was more rapidly turned over (degraded) at 78 weeks when collagen half-lives decreased. Together, this data implies age-specific collagen remodelling phases which may contribute to the pathogenesis of age-related kidney fibrosis.

### Distinct temporal patterns of matrix turnover during kidney ageing

To systematically characterise turnover dynamics, we performed hierarchical clustering of matrix protein half-lives throughout ageing, identifying distinct temporal patterns (**Figure 3A**). Six major clusters emerged: i) late-onset decrease (AGRN, TINAG, PZP, VTN, COL14A1), ii) early-onset decrease (LAMB2, LAMC1, COL4A3, COL4A1, CTSB, COL3A1, SERPINA3K, MFGE8), iii) transient early-onset decrease (COL6A1, COL6A3, SERPINA1A, COL6A2, TINAGL1, COL12A1, COL4A2, NID1, COL1A1, COL1A2, LAMA5), iv) steady increase (MEP1B, COL15A1, CTSD, ANXA5, CTSA, SERPINA1D), v) late-onset increase (FBN1, LAMB1, MEP1A, NID2, FGG, FN1, TGM2), and vi) transient early-onset increase (LGALS1, TGM1, FGA, FGB). Basement membrane components clustered predominantly into groups exhibiting transient or sustained early-onset decreases in turnover beginning by 52 weeks, consistent with early matrix stabilisation and decreased degradation in later life, providing an explanation for age-related GBM thickening and fibrosis. Interestingly, nidogen 1 and nidogen 2 displayed divergent turnover trajectories, with nidogen 1 showing progressive reduction in turnover from adulthood onward, whilst nidogen 2 demonstrated increased turnover with age, suggesting differential functional regulation within the basement membrane. Matrix proteins showing sustained turnover changes (steadily increasing or decreasing from 22 to 78 weeks) were particularly notable (**Figure 3B**). Proteins with sustained turnover increases included basement membrane associated collagen XV, proteases and matrix regulators. Conversely, proteins with sustained turnover decreases included basement membrane components along with network-forming and fibrillar collagens. This divergent regulation suggests distinct mechanisms controlling structural versus regulatory matrix components during ageing.

**Figure 3:**
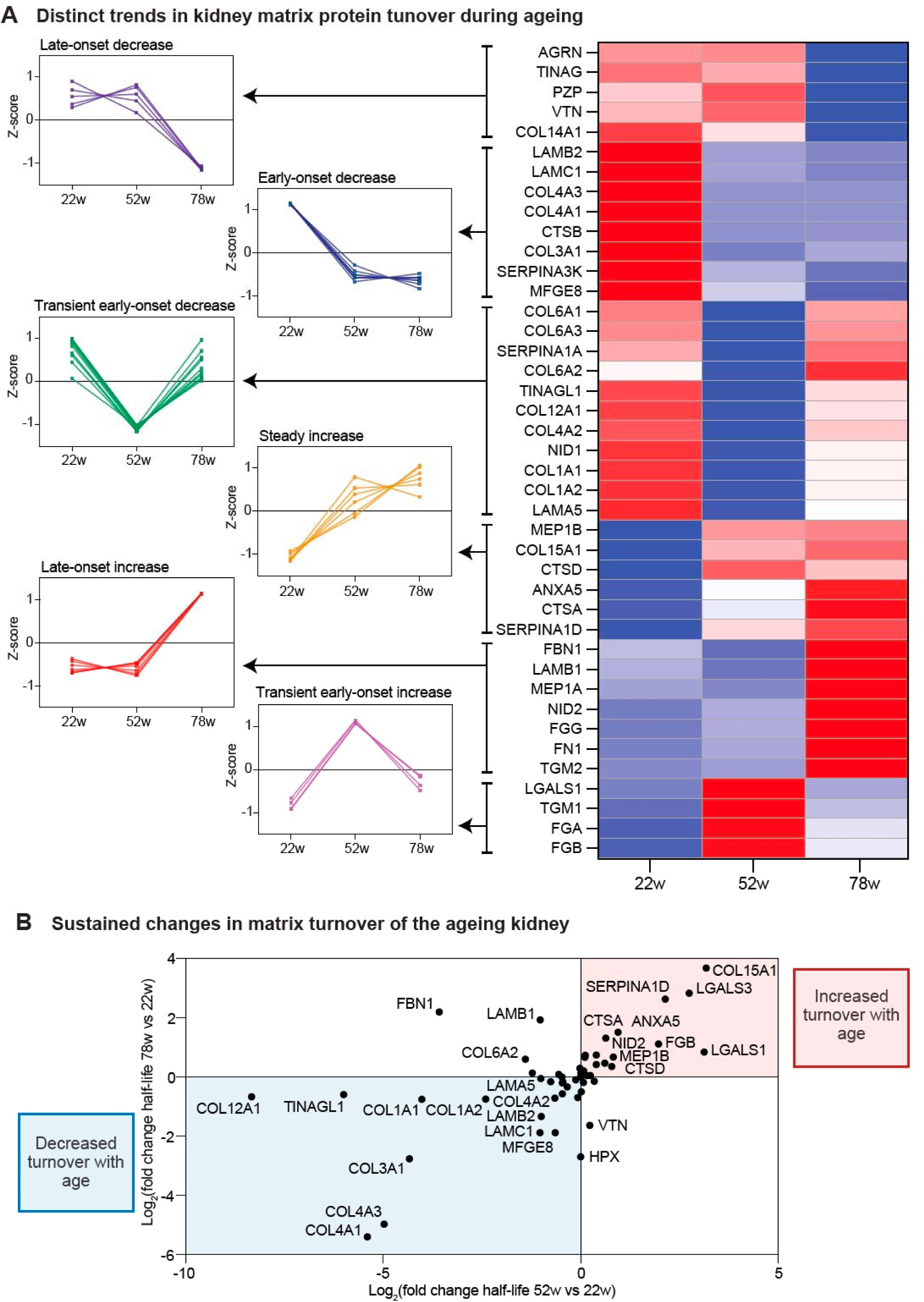
Distinct temporal patterns of matrix turnover during kidney ageing. **A**) Heat map of matrix protein z-scores calculated from heavy-to-light ratios from the matrix-enriched fraction isolated from 8-, 22-, 52-, and 78-week mice. Proteins were assigned into one of five groups following hierarchical clustering. **i)** Proteins in cluster 1 showed late-onset decrease in turnover. **ii)** Proteins in cluster 2 showed early-onset decrease in turnover. **iii)** Proteins in cluster 3 showed transient early-onset decrease in turnover. **iv**) Proteins in cluster 4 showed a steady increase in turnover. **v**) Proteins in cluster 5 showed late-onset increase in turnover. **vi**) Proteins in cluster 6 showed transient early-onset increase in turnover. **B**) Patterns of altered turnover in ageing comparing fold change in turnover (half-life) at 52-weeks and 78-weeks with 22-weeks demonstrating matrix proteins with sustained increase (shown in red) or sustained decrease (shown in blue) in turnover with advancing age. N=3 biological replicates per condition for all analysis.

### Predicted structural alteration in kidney matrix proteins

To gain mechanistic insight into kidney matrix proteolytic cleavage events and matrix reorganisation during development and ageing we used peptide location fingerprinting (PLF). Interpretation of PLF data is based on the relative accessibility of proteins to trypsin digestion, where increased peptide yield at a given time point is caused by increased accessibility to trypsin, and reduced peptide yield, decreased accessibility (**Figure 4A**). Changes in peptide yield therefore reflect differences in regional accessibility that may arise from proteolytic cleavage, higher-order conformational or network rearrangement, or accumulation of damage modifications which influence matrix reorganisation. Proteotypic peptides identified by MS/MS ion searches from the matrix-enriched fraction of kidneys from 8-, 52-, and 78-week mice were compared with those from adult 22-week mice. This analysis identified 77 kidney matrix proteins with statistically significant structure-associated differences (**Supplementary Figure 3A**). Of these, 45 matrix proteins demonstrated significant structural alteration relative to 22-week mice in both development (8 weeks) and ageing (52 and 78 weeks). The substantial overlap between developmental and ageing-associated changes represented 60% of structurally altered matrix proteins and notably, almost all core matrix proteins and basement membrane components had significant structural alteration during development and ageing. Based on convergent or divergent patterns of structural alteration during development and ageing, combined with their distinct turnover dynamics, we selected a focused subset of matrix proteins for further analysis. Meprin α (MEP1A) and β (MEP1B) were chosen as key matrix remodelling enzymes exhibiting age-dependent changes in turnover, implicating altered proteolytic activity with ageing. Basement membrane components nidogen 1 (NID1), nidogen 2 (NID2), and laminin-γ1 (LAMC1) were selected due to their sustained reductions in turnover and divergent regulation, highlighting potential changes in basement membrane stability and organisation. Collagen VI (COL6A1, COL6A2) and collagen XV (COL15A1) were chosen as representative interstitial and basement membrane associated collagens with dynamic and age-dependent turnover profiles, reflecting both developmental remodelling and age-associated matrix reorganisation.

**Figure 4:**
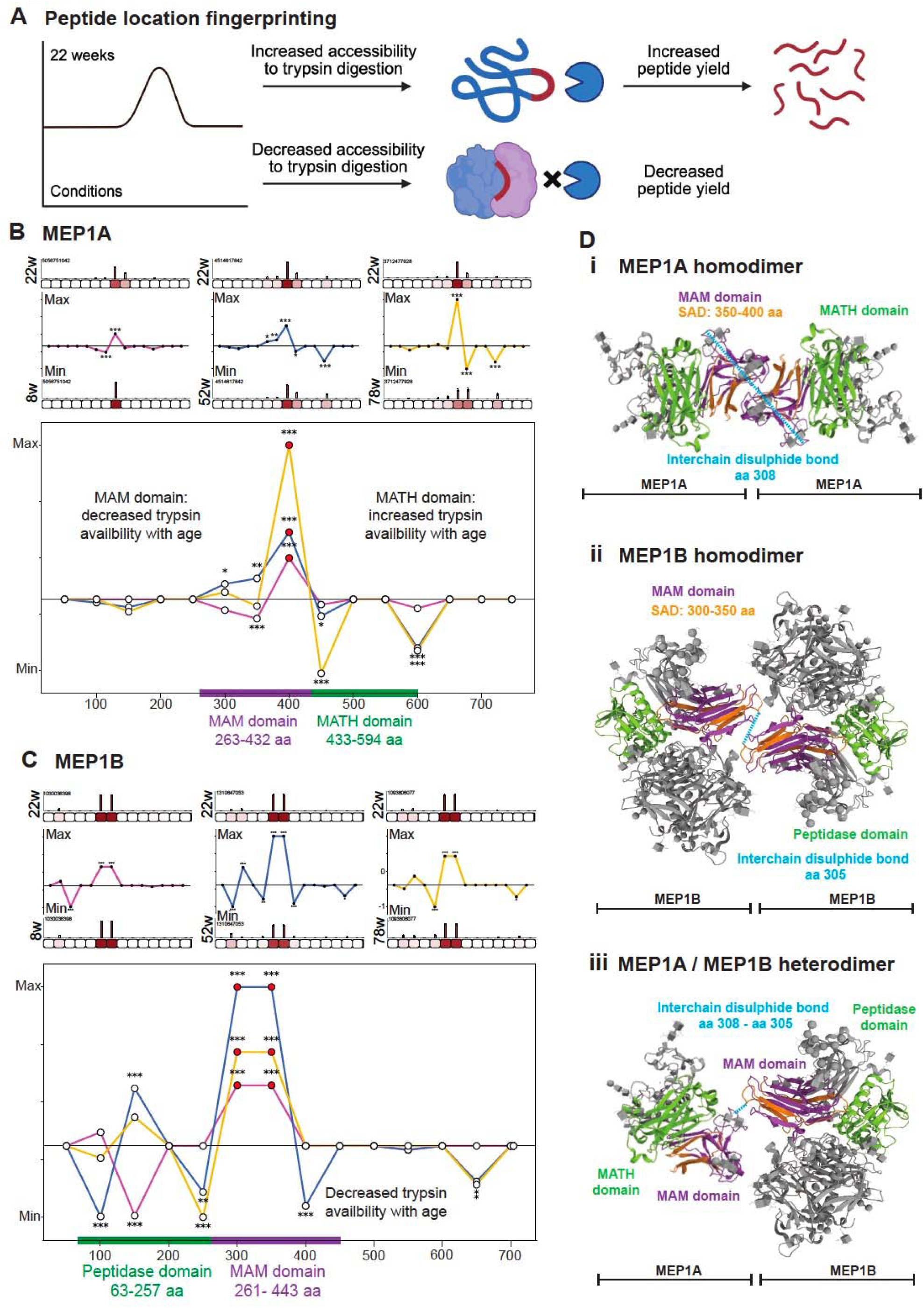
Meprin proteases display age-dependent structural changes within MAM domains. During PLF analysis, proteins were segmented into 50 amino acid (AA)-sized segments, proteotypic peptides were mapped and quantified (bar graphs = averaged, normalised summed intensities per segment, error bars = standard deviation) and statistically tested 8 weeks, 52 weeks and 78 weeks compared to 22 weeks (unpaired Bonferroni-corrected, repeated measures ANOVA; *p=<0.05, **p=<0.01, ***p=<0.001. Average peptide yields per segment in 8-weeks, 52-weeks and 78-weeks were then subtracted from 22-weeks and divided by segment length (50AA) to reveal differences along protein structure. Summed peptide intensities per segment were min/max normalised based on summed ion intensity to produce comparable line graphs along the protein structure (y axes = 22-8 weeks intensity/segment in magenta, 22-52 weeks intensity/segment length in blue, 22-78 weeks intensity/segment in yellow. For each analysis, structural differences are presented for 8- vs 22-week (left panel), 52- vs 22-week (middle panel), 78- vs 22-week (right panel) and all conditions (lower panel). **A**) Schematic demonstrating interpretation of PLF analysis; a peak in the direction of 22 week signifies increased accessibility to trypsin digestion (increased peptide yield) at 22-weeks compared to other conditions (8-, 52-, and 78-weeks), and decreased accessibility to trypsin digestion (decreased peptide yield) at 8-, 52-, and 78-weeks compared to 22-weeks. **B**) MEP1A demonstrated significantly higher peptide yield at 22-weeks compared to 8-, 52- and 78-weeks which corresponded with the MAM domain structure (shown by purple line), indicating decreased trypsin availability with age. Significantly altered peptide yield was also detected along the MATH domain (shown with green bar) at 52-, and 78-week mice compared to 22-weeks. **C**) MEP1B demonstrated significantly higher peptide yield at 22-weeks compared to 8-, 52-, and 78- weeks which corresponded to the MAM domain (shown by purple bar), indicating decreased trypsin availability with age. Significantly altered peptide yield was also detected along the peptidase domain (shown by green bar) at 8-, 52-, and 78-week mice compared to 22-weeks. **D) i,** Crystal structure of MEP1A presented in a homodimerized structure with MAM domains (purple) interacting in an inaccessible confirmation representing predicted MEP1A structure at 8-, 52-, and 78-weeks compared to 22-weeks. Region of SAD (structure-associated differences) detected by PLF analysis is shown in orange. PDB: 7UAC https://doi.org/10.1038/s41467-022-33893-7). **ii**, Crystal structure of MEP1B presented in a homodimerized structure with MAM domains interacting in inaccessible confirmation and peptidase domain in an accessible confirmation inferring active protease activity, representing predicted MEP1B structure at 8-, 52-, and 78-weeks compared to 22-weeks. Region of SAD (structure-associated differences) detected by PLF analysis is shown in orange. PDB: 4GWN https://doi.org/10.1073/pnas.1211076109). **iii,** Crystal structure of MEP1A and MEP1B presented in heterodimerized structure with MAM domains (shown in purple) interacting in an inaccessible conformation, and MATH and peptidase domains (shown in green) in an accessible confirmation representing predicted MEP1A/MEB1B heterodimer structure and inferring active protease activity at 8-, 52-, and 78-weeks compared to 22-weeks. N=3 biological replicates per condition for all analysis.

### Predicted age-dependent structural changes in meprin proteases

Meprins are multidomain oligomeric metalloproteinases involved in inflammation, fibrosis and degradation of the extracellular matrix [68–71]. The meprin family comprises two homologous subunits, MEP1A and MEP1B, which form disulphide-linked homo-and hetero-oligomers at the cell surface and can be released into the extracellular space. MEP1A and MEP1B have similar domain architecture and share MAM domains which are important in protein-protein interactions and oligomerisation [72–74]. With PLF we predicted age-dependent structural alterations in both MEP1A and MEP1B, consistent with altered protease activity during kidney development and ageing (**Figure 4B and 4C**). For MEP1A, altered trypsin accessibility localised to the MAM domain (aa 263-432) and adjacent MATH domain (aa 433-594) when comparing developing (8 weeks) and ageing (52 and 78 weeks) kidneys with adult controls (22 weeks) (**Figure 4B**). These domains mediate oligomer assembly required for full proteolytic activity [72, 73, 75]. Notably, PLF revealed reduced trypsin accessibility within the MAM domain in ageing alongside increased accessibility within the MATH domain. This reciprocal pattern is consistent with MAM-mediated homodimer assembly, in which MAM domains are buried within inter-subunit interfaces, whilst MATH domains become more exposed (**Figure 4D**). These changes support age-associated rearrangement of MEP1A homo-oligomeric architecture, with potential consequences for protease accessibility and matrix remodelling capacity. Our PLF analysis of MEP1B similarly predicted age-dependent structural alterations, with altered peptidase domain (aa 63–257) and reduced MAM domain (aa 261-443) trypsin accessibility when comparing developing (8 weeks) and ageing (52 and 78 weeks) kidneys to adult controls (22 weeks) (**Figure 4C**). The peptidase domain contains the catalytic zinc binding-site, which must be structurally exposed for substrate binding [76]. Structural studies indicate MAM domains contribute to protease organisation within active complexes [74, 77], placing these domains near dimerisation interfaces whilst positioning the peptidase domain for substrate access (**Figure 4D**). Decreased MAM domain accessibility during development and ageing is consistent with engagement in dimeric conformations, whilst altered peptidase domain accessibility predicts changes in catalytic activity. Shedding of membrane-bound meprin β, by ADAM10 (a disintegrin and metalloproteinase) allows for its activation and release [78, 79], increasing proteolytic activity. Interestingly, MEP1B was significantly increased in abundance in young mice, and ADAM10 was detected as structurally altered at this time-point (**Supplementary Figure 3A**). Additionally, increased peptidase domain accessibility during advanced age coincided with significantly increased MEP1B turnover, highlighting a potential role for MEP1B-mediated proteolytic remodelling in the ageing kidney.

### Predicted structural alterations in core basement membrane components

The mature GBM comprises four major macromolecules; type IV collagen, laminin, nidogen and perlecan [80] (**Figure 5A**). Nidogen 1 and 2 (NID1, NID2) are basement membrane bridging proteins that stabilise the GBM by linking laminin-521 and collagen IV networks through distinct G2 and G3 domains [81–83]. The G2 domain binds perlecan and collagen IV [81, 84, 85], whilst the G3 domain binds the laminin γ-subunit short arm [86] (**Figure 5A**). Since NID1 and NID2 had opposing turnover profiles with NID 1 showing significantly decreased turnover throughout ageing (similar to LAMC1), whilst NID2 turnover increased at 78 weeks we used PLF to assess structure associated difference in nidogens. We found that NID1 had domain-specific, age-dependent changes in trypsin accessibility (**Figure 5B**). There was significantly higher peptide yield at 22 weeks compared to 52 and 78 weeks within the G2 domain suggesting reduced trypsin accessibility with age. Conversely, we found significantly lower peptide yield within the G3 domain at 22 weeks compared with 8, 52 and 78 weeks suggesting increased trypsin accessibility during development and ageing. These reciprocal changes imply age-associated changes in the interaction state of NID1, which could affect stabilisation of both collagen IV–perlecan and laminin 521 networks. Our PLF analysis of NID2 demonstrated a pattern distinct from NID1 (**Figure 5C**). Significantly lower peptide yield at 22 weeks compared to 52 and 78 weeks within the G2 domain suggests increased trypsin accessibility with age. Additionally, altered peptide yield across the G3 domain at 8, 52 and 78 weeks relative to 22 weeks suggests dynamic age-dependent changes in NID2 laminin-binding across development and ageing. Overall, these findings suggest differential interaction states of nidogen isoforms during kidney ageing.

**Figure 5:**
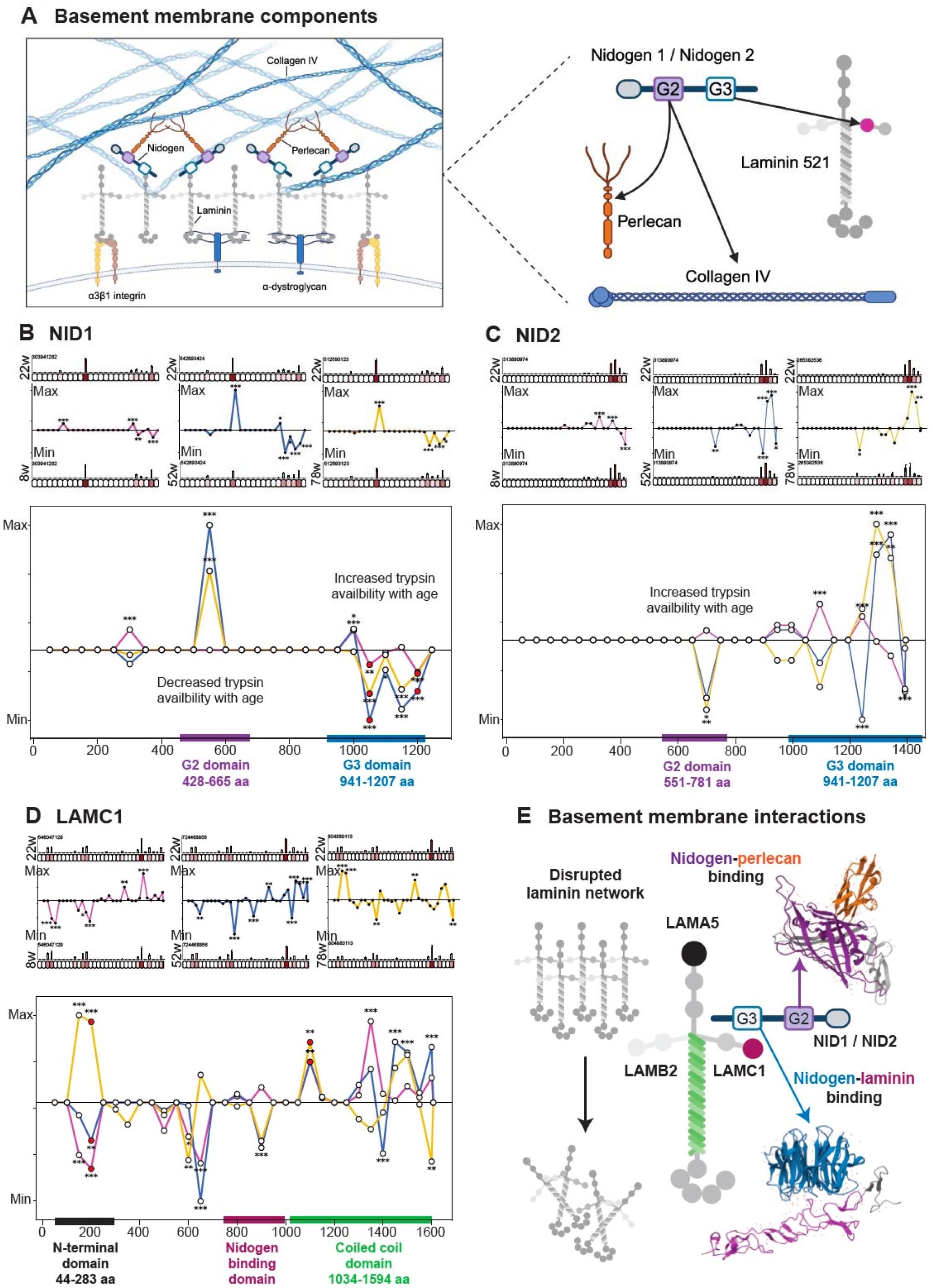
Basement membrane core components display structural alterations within regions of higher-order interaction. PLF analysis was completed as described in Figure 4. **A**) Left panel: schematic visualising the core components of glomerular basement membrane including collagen IV, laminin 521, nidogen and perlecan. Right panel: schematic demonstrating basement membrane component interactions; nidogen G2 domain binds perlecan and collagen IV whilst nidogen G3 domain binds laminin-γ1 within the laminin 521 network. Created with BioRender.com. **B**) NID1 demonstrated significantly higher peptide yield at 22-weeks compared to 52- and 78-weeks which corresponded with the G2 domain (shown by purple bar), indicating decreased trypsin availability with age. Significantly lower peptide yield was detected along the G3 domain (shown with blue bar) at 22-weeks compared to 8, 52-, and 78-week mice, indicating increased trypsin availability with age. **C**) NID2 demonstrated significantly lower peptide yield at 22-weeks compared to 52- and 78-weeks which corresponded with the G2 domain (shown by purple bar), indicating increased trypsin availability with age. Significantly altered peptide yield was also detected along the G3 domain (shown by blue bar) at 8, 52-, and 78-week mice compared to 22-weeks. **D**) LAMC1 demonstrated widespread structure-associated differences throughout the protein sequence at 8-, 52-, and 78-weeks compared to 22-weeks. This included altered peptide yield detected in the N-terminal domain (shown by black bar), the nidogen-binding domain (shown by magenta bar), and the coiled-coil domain (shown by green bar) and infers alterations in higher-order structure of the laminin 521 network in ageing. **E**) Schematic of significantly altered binding domains in basement membrane components across the kidney life course showing the crystal structure of nidogen-perlecan binding, mediated through the nidogen G2 domain (PDB: 1GL4), the crystal structure of nidogen-laminin-γ1 binding, mediated through the nidogen G3 domain (PDB: 1NPE), and predicted disruption in the laminin 521 network caused by alteration in higher order structure during ageing. N=3 biological replicates per condition for all analysis.

Our PLF analysis of LAMC1 revealed extensive structural differences at 8, 52 and 78 weeks compared to 22 weeks (**Figure 5D**). There was altered peptide yield within the N-terminal domain, nidogen-binding domain, and coiled-coil domain, suggesting age-dependent changes in the higher-order structure of LAMC1. These changes could affect laminin network stability and may represent damage accumulation. Collectively, this data indicates coordinated, age-dependent structural reorganisation of nidogen-laminin and nidogen-perlecan interactions, alongside widespread structural alterations affecting laminin-521 network organisation (**Figure 5E**). Given these basement membrane alterations, we next examined whether matrix proteins involved in microfibrillar assembly and cell–matrix interactions also exhibit structural changes during ageing, and we applied PLF to collagen VI and collagen XV.

### Predicted domain-specific structural alterations in collagen VI

Using PLF we identified significant, age-dependent differences in collagen VI peptide yield when comparing adult kidneys (22 weeks) with developing (8 weeks) and ageing (52 and 78 weeks) time points (**Figure 6A and 6B**). Differences in COL6A1 localised predominantly to the N-terminal von Willebrand factor A (VWFA1) domain and the C-terminal VWFA2/3 domains (**Figure 6A**), which are required for monomer assembly and microfibril formation. Collagen VI monomers assemble into antiparallel dimers through N- and C-terminal interactions, which then form tetramers that associate end-to-end to generate microfibrils [87] (**Figure 6C**). Altered peptide yield within these assembly domains indicates age-dependent changes in domain accessibility, consistent with supramolecular reorganisation of collagen VI assemblies during development and ageing. Similarly, COL6A2 demonstrated significant age-associated differences within the C-terminal VWFA2/3 domains (**Figure 6B**), supporting remodelling across constituent chains. In addition, we found significantly increased peptide yield at 22 weeks compared to 52 and 78 weeks within the RGD-containing domain, suggesting reduced trypsin accessibility with ageing. The RGD motif, located within the triple helical region, mediates integrin-dependent cell binding [88, 89] (**Figure 6C**) and reduced accessibility of this domain suggests age-dependent structural reorganisation that may alter the exposure of cell surface protein interaction sites in collagen VI microfibrils.

**Figure 6:**
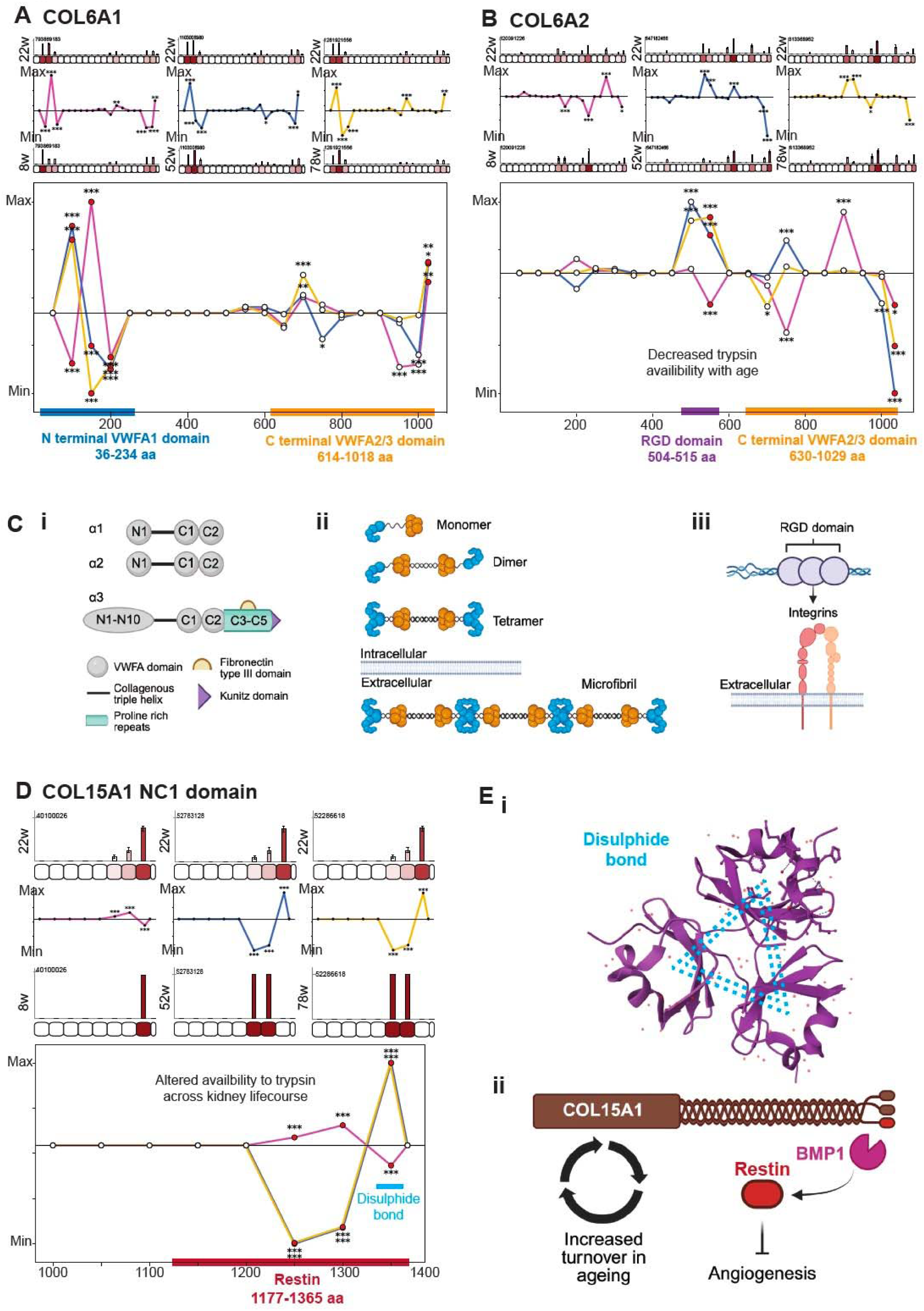
Domain-specific structural alterations in collagen VI. PLF analysis was completed as described in Figure 4. **A**) COL6A1 demonstrated significantly altered peptide yield at 22-weeks compared to 8-, 52- and 78-weeks in the N terminal VWFA (Von Willibrand Factor A) 1 domain (shown by blue bar) and C terminal VWFA2/3 domains (shown by orange bar). **B**) COL6A2 demonstrated significantly altered peptide yield at 22-weeks compared to 8-, 52- and 78-weeks in the C terminal VWFA2/3 domains (shown by orange bar). Significantly increased peptide yield was detected at 22-weeks compared to 52-, and 78-weeks in the COL6A2 RGD domain (shown by purple bar), indicating decreased trypsin availability in ageing and inferring structural instability of collagen IV tetramers, exposing RGD domains for increased cell binding. **C**) Schematic illustrating collagen VI structure, microfibril formation and cell binding. **i**) Domain arrangement of the three most common collagen VI α chains (α1-3) showing the C- and N-terminal VWFA domains which are required for higher-order assembly. **ii**) Collagen VI heterotrimeric monomers form from one α1, one α2 and one long α-chain (or α3-6) chain. Triple-helical monomers form disulfide-linked anti-parallel dimers which in turn associate to form tetramers, which are secreted into the extracellular space where microfibrils are formed by interlinking tetramers. The C-terminal globular regions are shown in orange, and the N-terminal globular domains are shown in blue, correlating to regions of significant structure—associated differences detected in COL6A1 and COL6A2 by PLF analysis. **iii**) Schematic illustrating an accessible RGD sequence in structurally disrupted collagen VI microfibril, enabling cell-interaction through integrin-binding in ageing. **D**) COL15A1 demonstrated widespread structure-associated differences throughout the NC1 domain, corresponding to the matrikine restin, indicating alterations in restin production during the kidney lifespan. Significantly higher peptide yield was detected at 22-weeks compared to 52-, and 78-weeks, mapping to a disulphide bond within the NC1 domain, whilst significantly lower peptide yield was detected in this region in 22-week compared to 8-week mice. **E**) **i**) Schematic demonstrating NC1 COL15A1 trimerization forming a stable COL15 protein, mapping to the region of structural associated differences detected by PLF analysis (PDB 3N3F). **ii**) Schematic demonstrating relationship between COL15A1 turnover, which increased in ageing, and predicted alteration in restin matrikine cleavage which mediates anti-angiogenic effects. N=3 biological replicates per condition for all analysis.

### Predicted age-associated matrikine release from collagen XV

Finally we performed PLF analysis of COL15A1 and found widespread structure-associated differences throughout the NC1 domain, which encodes the anti-angiogenic matrikine restin [27] (**Figure 6D**). Here, significantly lower peptide yield at 22 weeks compared to 52 and 78 weeks, indicates increased trypsin accessibility in ageing (52 and 78 weeks) compared to 22 weeks and the opposite was observed for young (8 week) mice. However, significantly higher peptide yield at 22 weeks compared to 52 and 78 weeks was detected within regions corresponding to a disulphide bond involved in NC1 domain stabilisation indicating reduced trypsin accessibility with ageing. The NC1 domain is required for collagen XV trimerization [90], and following basement membrane incorporation, can be proteolytically cleaved to release the bioactive restin fragment [27] (**Figure 6E**). These data suggest age-dependent structural alterations may influence both NC1 domain stability and proteolytic accessibility, potentially affecting restin generation. Soluble restin would likely yield more peptides than intact collagen XV, so higher peptide yields in this region may indicate increasing restin levels with age. Notably, collagen XV demonstrated sustained turnover increases at both 52 and 78 weeks, indicating ongoing remodelling during ageing. This occurred alongside reduced abundance in young (8-week) kidneys and increased abundance at 52 weeks, consistent with dynamic regulation across the kidney lifespan. The combination of sustained turnover, increased abundance, and altered NC1 domain accessibility may have functional consequences for angiogenic regulation in the ageing kidney.

## Discussion

Progressive accumulation of extracellular matrix is a hallmark of kidney fibrosis and age-related decline in kidney function [16], yet the dynamic processes governing matrix remodelling across the physiological lifespan remain poorly understood. In this study, we conducted a systems level investigation of kidney matrix homeostasis by integrating heavy-isotope labelled proteomics with the prediction tool PLF to define the natural dynamics of protein turnover and higher order structural organisation of the kidney matrix from development to advanced age. We discovered changes in matrix composition and turnover across the lifespan indicating active assembly during development, homeostatic maintenance in adulthood, and progressive dysregulation with ageing. Furthermore, we identified specific patterns of altered turnover for proteases, basement membrane components, and non-fibrillar collagens that predict disrupted proteolytic processing, providing mechanistic insight into the pro-fibrotic predisposition of the ageing kidney.

For the first time, we have defined the natural dynamics of protein turnover in the kidney matrix across four timepoints of the murine lifespan, ranging from young (8 weeks), adult (22 weeks), aged (52 weeks), and advanced aged (78 weeks). In young mice, kidney matrix proteins displayed significantly shorter half-lives compared to adult mice, indicating markedly accelerated turnover during post-natal kidney maturation. By adulthood, matrix protein turnover had slowed substantially, indicating establishment of more stable, homeostatic extracellular matrix. Such dynamics are consistent with studies demonstrating progressive basement membrane assembly from embryonic stages to early postnatal life [39], with our data revealing this process extends into early adulthood before achieving steady-state dynamics.

Beyond adulthood, kidney matrix composition and turnover underwent progressive changes indicative of disrupted homeostasis. Previous mass spectrometry-based proteomics studies have described a signature of altered matrix in human kidney ageing, demonstrating age-associated increases in structural components and matrix regulators alongside decreases in basement membrane components [38]. Our data aligns, demonstrating sustained increases in collagen VI chains and matrix regulators with similar patterns in basement membrane components (decreases in collagen IV, LAMB2, NID2, HSPG2, TINAG and increases in NID1, LAMB1 and LAMA4). Our complimentary findings imply that the loss of equilibrium between matrix degradation and synthesis is conserved across species and the addition of turnover data in our study provides mechanistic insight. Basement membrane components including collagen IV α-chains, laminin-521 chains, perlecan, and nidogens demonstrated sustained decreases in turnover from development throughout ageing, with collagen IV half-lives increasing from weeks in young mice, to months in adult mice, before extending to years in aged mice (52 and 78 weeks). Previous turnover studies in *Drosophila* [48] and *C. elegans* [49] demonstrates basement membrane components nidogen and perlecan to have extremely short half-lives, in the order of hours, and revealed high mobility of these components within more static laminin and collagen IV scaffolds. Until now, half-lives of kidney BM components in mammalian systems have not been quantified and we now confirm a dynamic property of nidogens and perlecan, with half-lives ranging from days to weeks, compared to much longer-lived laminins and collagen IV chains. Furthermore, PLF analysis identified structural alterations within nidogen and laminin-y1 binding interfaces, predicting weakened nidogen-laminin-collagen IV interactions and progressive basement membrane destabilisation with age. As basement membrane components become increasingly long-lived over the lifespan, accumulation of damage modifications likely compromises basement membrane mechanical properties and permeability, contributing to proteinuria and basement membrane thickening characteristic of age-related kidney disease.

Our integration of structural, abundance and turnover data revealed dynamic collagen VI remodelling across the kidney lifespan. Collagen VI is markedly upregulated in kidney disease [91–93], however little is known about its role in physiological kidney ageing. We demonstrate increased abundance of all collagen VI chains (COL6A1, COL6A2, COL6A3) at 52 weeks that was not sustained at 78 weeks. Turnover rates dramatically decreased at 52 weeks, but increased again at 78 weeks, suggesting progressive accumulation of long-lived microfibrils occurs early in age-related kidney fibrosis with subsequent degradation in advanced age. The C-terminal of collagen VI-α3 is cleaved to release the pro-fibrotic matrikine endotrophin [30] and high serum levels correlate with disease progression and degree of kidney fibrosis in CKD [93–95]. Whilst our PLF analysis was unable to quantify structural alterations along COL6A3, widespread structural differences were detected throughout the VWFA domains of COL6A1 and COL6A2, indicating altered monomer assembly and microfibril formation. We propose that endotrophin may serve as a biomarker for kidney fibrosis associated with ageing, reflecting degradation of structurally altered collagen VI microfibrils in advanced age.

Collagen XV has been identified as part of the consistent matrix signature that increases across kidney ageing and disease [38] and deposition has been observed in kidney fibrosis [25]. Whilst our data supports increased collagen XV during ageing, we demonstrate a distinct turnover profile compared to other interstitial collagens and basement membrane components. We observed sustained rapid turnover (half-live of 7-9 days) during ageing despite increased abundance, contrasting sharply with the dramatic stabilisation observed in collagens I, III, IV and VI (half-lives > 400 days). Collagen XV contains an anti-angiogenic restin domain, a matrikine that upon cleavage inhibits angiogenesis through modulation of endothelial cell migration [27, 30]. Endostatin, a matrikine which shares sequence homology with restin upon cleavage from collagen XVIII, demonstrates increased abundance in aged mice [96] and its overexpression induces interstitial fibrosis [97]. Here, we show age-dependent structural alteration throughout the collagen XV NC1 domain, corresponding to restin, suggesting this related matrikine may also play a bioactive role in kidney fibrosis associated with ageing. The paradoxical pattern of increased abundance with maintained turnover suggests that collagen XV accumulation reflects active, ongoing matrix remodelling rather than impaired degradation, potentially indicating altered proteolytic processing as the primary mechanism driving its contribution to the fibrotic phenotype.

Upregulation of collagens I and III is widely accepted as a major contributor to extracellular matrix accumulation in kidney fibrosis [65, 98]. Recent proteomic comparisons found no significant differential abundance of COL1A1, COL1A2 and COL3A1 in human foetal versus mature kidney [32] or in human ageing kidney [38], although trending increases were observed in the oldest age group (69 years). However, in the context of kidney disease, there was consistent increases in collagen I and III across mouse models of glomerular disease and multiple human kidney diseases [38]. In our mouse ageing study (8-78 weeks, approximately equivalent to human ageing up to 60 years), we observed decreased collagen I and III abundance throughout ageing, yet the most striking finding was dramatic changes in turnover dynamics with half-lives increasing from 10-22 days in young mice to over 400 days in aged mice (52 weeks) which represents a more than 40-fold decrease in turnover, suggesting a near absence of degradation. This pattern suggests that collagen accumulation in physiological aging may be a late phenomenon, potentially emerging only at advanced age (beyond 78 weeks in mice, equivalent to >60-70 years in humans), after progressive loss of remodelling capacity has reached a critical threshold. Indeed, a recent urinary peptidomics study demonstrated abundance of COL1A1 peptides decreased with increasing with age [99], supporting our observation that age-related kidney fibrosis results primarily from attenuated collagen degradation rather than increased synthesis. Interestingly, at our oldest time-point (78 weeks), collagen (type I, III and VI) half-lives began to decrease in parallel with increased turnover of collagen cross-linking enzyme TGM2, suggesting cross-linking pathways are diminished in advanced age and supporting the role of TGM2 in the progression of kidney fibrosis [67, 97, 100].

Decorin (DCN), a proteoglycan which exerts antifibrotic effects by inhibiting TGF-β and matrix accumulation [101], was notably decreased in both 52-week and 78-week mice. TGF-β signalling contributes to fibrosis development [102], causing collagen accumulation, cross-linking and suppression of collagen degradation [103, 104]. The combination of decreased decorin alongside increased collagen VI accumulation and dramatically stabilised fibrillar collagens, creates a pro-fibrotic environment in which normal protective mechanisms against matrix accumulation are compromised. Our findings suggest that decorin replacement could be a preventative strategy in kidney ageing, particularly given that established therapeutic approaches have shown efficacy in other fibrotic conditions [64].

The metalloproteases, meprin α and meprin β (MEP1A, MEP1B) have pro-inflammatory activity and contribute to extracellular remodelling in fibrosis [68, 69]. A well-described physiological function of meprins is maturation of fibrillar procollagens I and III through cleavage of N- and C-terminal prodomains [105, 106], a prerequisite for collagen fibril assembly. Meprins also show substrate specificity for basement membrane components collagen IV and nidogen 1 [68, 107], cleavage of which generates pro-inflammatory matrikines. Interestingly, in a mouse model of kidney injury, meprin inhibition correlated with decreased excretion of fragmented nidogen 1 into the urine [107]. Our data demonstrates complex regulation of meprins across the lifespan. MEP1B abundance was elevated during development alongside rapid basement membrane turnover. In murine kidneys, meprin α can either associate with membrane-bound meprin β or self-associate to form homo-oligomers of meprin α subunits, which are secreted [73, 108]. The ectodomain of meprin β can be shed by ADAM10 [78], suggesting ADAM10 regulates proteolytic action of membrane bound meprins. Interestingly, ADAM10 was one of the only matrix proteins detected by PLF analysis to display structural alteration in young mice alone, supporting its role in early kidney matrix remodelling during development. During early ageing (52 weeks), both MEP1A and MEP1B increased in abundance, with MEP1A showing late onset increases in turnover and MEP1B showing steady increases throughout ageing. This increase in meprin synthesis coincided with significantly decreased turnover of fibrillar collagens, suggesting meprin-mediated collagen stabilisation as a mechanism in physiological ageing. Additionally, age-dependent structural alterations were detected in nidogen isoforms along with late-onset increases in their turnover. PLF revealed age-dependant structural alterations in meprin MAM and MATH domains, suggesting shifts in oligomerisation that may favour active enzyme conformation and proteolytic efficiency. Collectively, our data supports meprins as key regulators of age-related kidney matrix remodelling and basement membrane proteolysis, highlighting their potential for therapeutic targeting.

Overall, we provide the first comprehensive study of kidney matrix protein turnover across mammalian lifespan, revealing that age-related fibrosis is characterised by progressive loss of matrix remodelling capacity rather than increased synthesis. The dramatic stabilisation of fibrillar and basement membrane collagens, alongside suppression of the anti-fibrotic proteoglycan decorin, creates a pro-fibrotic state that predisposes the ageing kidney to injury. Integration of turnover data with peptide location fingerprinting predicts altered protease activity, basement membrane stability, and collagen assembly with potential implications for matrikine biology. The quantification of matrix protein half-lives across the lifespan provides essential baseline data for future interventional studies aimed at understanding the mechanistic basis of age-related kidney fibrosis and developing strategies to preserve matrix homeostasis.

## Materials and methods

### Animals

The care and use of all mice in this study was carried out in accordance with the UK Home Office regulations, UK Animals (Scientific Procedures) Act of 1986 under the Home Office Project License’s: P1AE9A736, PP4262564, PP708858 and LO45CA465. Mice were maintained in 12-h light/12-h dark cycle at 20°C to 22°C, humidity of 40-50%, with *ad libitum* access to water. Wildtype (C57BL/6J) male mice were fed “light diet” (standard rodent chow) or “heavy diet”; Mouse Express L-Lysine (^13^C_6_, 99%; MLK-LYS-C) purchased from CK Isotopes Ltd (UK), as per the labelling experimental design. All mice gained weight in accordance with reference laboratory data for C57BL/6J mice.

### Preparation of tissues

Mice were sacrificed by cervical neck dislocation and kidneys were immediately dissected. The surrounding fat, adrenal glands and renal capsule were removed, and kidneys for downstream proteomic experiments were snap frozen and stored at −80°C.

### Pulsed ^13^C-Lys labelling experimental design

Wildtype (C57BL/6J) male mice (n=3 per age group) were pulse-labelled with ^13^C-Lys (heavy diet) for 4 weeks before sacrifice for tissue harvesting at either 8 weeks (young mice), 22 weeks (adult mice), 52 weeks (old mice) or 78 weeks (aged mice) (**see Figure 1A**). The purpose of pulse-labelling for different periods was to monitor label incorporation over time to allow protein half-life calculation using a recognised mass spectrometry workflow [66].

### Matrix-enrichment protocol for proteomic experiments

Frozen-thawed kidneys were placed in a glass dish, cut into small pieces (<1 mm^3^) and washed three times with ice-cold PBS. To solubilise the kidney, 200 μl of SL-DOC buffer (1.1% (vol/vol) sodium dodecanoate; 0.3% (vol/vol) sodium deoxycholate; 25 mM ammonium bicarbonate; 0.5 mM dithiothreitol (DTT); protease and phosphate inhibitors as per manufacturer’s guidance (Sigma); pH 7) was added to the kidney tissue along with six 1.6 mm steel beads. Samples were homogenised at maximum speed for 5 minutes at 4°C using a Bullet Blender Tissue Homogeniser (Next Advance, New York, USA). Samples were alkylated with IAA (final concentration, 45 mM), before quenching with DTT. Following centrifugation, supernatants were collected and transferred to LoBind Protein tubes. This sample was designated soluble fraction 1 (F1; cellular compartment). Remaining pellets were further solubilised using 75 μl of High Salt Buffer (3 M sodium chloride; 25 mM DTT; 25 mM ammonium bicarbonate, protease and phosphate inhibitors as per manufacturer’s guidance; pH 7) and further homogenised using a bullet blender at maximum speed for an additional 5 minutes. Following centrifugation, remaining pellets were sonicated in 300 μl of Lysis buffer (5% sodium dodecyl sulphate (Sigma); 50 mM triethylammonium bicarbonate (TEAB); pH 7), supplemented with protease and phosphatase inhibitors, using a Covaris LE220+ Sonicator (40W programme; 100 cycle per burst; duty factor 40%; peak incident power 500W). The protein samples were reduced with DTT (final concentration, 5 mM), followed by alkylation and quenching as above. To generate the matrix-enriched fraction 2 (F2; matrix compartment), the supernatants from the High Salt and Lysis buffer extractions were pooled. Protein concentrations in F1 and F2 were determined by BCA.

### Preparation of samples for mass spectrometry analysis

50 µg of protein from each sample was prepared in 5% SDS to a total volume of 50 µL and then acidified using 5 µL of 12% H_3_PO_4_. 350 µL binding buffer (90% methanol, 10% distilled water; 100 mM TEAB) was added and the samples loaded onto an S-Trap plate (ProtiFi) according to the manufacturer’s protocol. Bound proteins were washed in binding buffer and methyl tert-butyl ether (MTBE) and then digested overnight at 37°C with 5 µg of trypsin in Digestion buffer (50 mM TEAB; 0.1% formic acid, FA; at pH 8.5). Peptides were eluted from the column in 65 µL Digestion buffer, 65 µL 0.1% FA in distilled water, and finally with 30 µL 0.1% FA, 30% acetonitrile in water. Peptides were desalted using Oligo R3 resin beads, according to the manufacturer’s protocol, in a 96-well, 0.2 µm polyvinylidene fluoride filter plate. The immobilised peptides were washed twice with 0.1% FA prior to elution with 0.1% FA with 30% ACN, and lyophilisation.

### Mass spectrometry data acquisition

Peptides were resuspended in 10 µl 0.1% FA in 5% ACN. Liquid chromatography (LC) separation was performed on a Thermo RSLC system consisting of an NCP3200RS nano pump, WPS3000TPS autosampler and TCC3000RS column oven configured with buffer A as 0.1% (vol/vol) FA in water and buffer B as 0.1% (vol/vol) DA in ACN. The analytical column (Waters nanoEase M/Z Peptide CSH C18 Column, 130Å, 1.7µm, 75µm x 250mm) was kept at 35°C and at a flow rate of 300 nL/minute for 8 minutes. The injection valve was set to load before a separation consisting of a 105-minute multistage gradient ranging from 2-65% of buffer B. The LC system was coupled to a Thermo Exploris 480 mass spectrometry system via a Thermo Nanospray Flex ion source. The nanospray voltage was set at 1900V and the ion transfer tube temperature set to 275°C. Data was acquired in a data-dependent manner using a fixed cycle time of 2 seconds, an expected peak width of 15 seconds and a default charge state of +2. Full mass spectra were acquired in positive mode over a scan range of 300 to 1750Th, with a resolution of 120000, a normalised automatic gain control (AGC) target of 300% and a maximum fill time of 25 ms for a single microscan. Fragmentation data was obtained from signals with a charge state of +2 or +3 with an intensity over 5000 and they were dynamically excluded from further analysis for a period of 15 seconds after a single acquisition within a 10-ppm window. Fragmentation spectra were acquired with a resolution of 15000, a normalised collision energy of 30%, an AGC target of 300%, a first mass of 110Th, and a maximum fill time of 25 ms for a single microscan. All data was collected in profile mode.

### Proteomic data analysis

Mass spectrometry analysis was carried out using a SILAC 1plex (Lys6) method in Proteome Discoverer (version 3.2) (Thermo Fisher Scientific). Carbamidomethylation of cysteine was selected as a fixed modification and variable modifications included oxidation of methionine, hydroxylation of proline and phosphorylation of threonine, serine, and tyrosine. A maximum of 2 missed cleavages per peptide was allowed. The minimum precursor mass was set to 350 Da with a maximum of 5000 Da. Precursor mass tolerance was set to 10 ppm, fragment mass tolerance was 0.02 Da and minimum peptide length was 6. Peptides were searched against the mouse Swiss-Prot database with additional TreEMBL matrix protein sequences (unreviewed) using the SEQUEST HT search tool. Identified proteins were required to have a minimum false discovery rate (FDR) of 1% and at least 2 unique peptides. Known contaminants were removed. Abundance was calculated from the MS1 ion current peaks and combined for both heavy and light channels. Values were exported to Perseus (version 2.0.11.0) for abundance analysis, proteins were filtered for those with at least 3 calculated abundances, normalised to all proteins by median subtraction (either to all proteins or, for F2, to matrisome proteins) and then missing data was imputed based on standard deviation. Heavy-to-light ratios were determined using the raw abundance of a protein detected in the heavy channel, containing ^13^C-lysine, compared to the light channel for each repeat independently. Heavy-to-light ratios were converted to half-lives using published mass-spectrometry workflow developed by Alevra *et al* [66], with a minimum of 5 data points required for half-life calculation. Default parameters were used for modelling lysine pools and model fit was based on F1 samples from a pulse-time experiment where 22-week wildtype (C57BL/6) mice pulse-labelled with a ^13^C-Lys heavy diet for 1-, 2-, 4-, or 8-weeks (n=3 biological replicates per time-point). The purpose of using multiple labelling intervals was to generate sufficient data for modelling of protein half-lives. Protein and peptide lists were filtered for matrix proteins using R code (v4.1.2) based on their presence in MatrisomeDB [13]. Heatmaps and associated hierarchical clustering were generated using Instant Clue (v0.12.1). Z scores were calculated from average heavy-to-light ratios across repeats, and the mean and standard deviation were calculated across ages.

### Peptide location fingerprinting

*We used* our previously developed webtool as previously described [59–61]. (https://www.manchesterproteome.manchester.ac.uk/#/MPLF). Peptides (FDR threshold <0.01) were imported into the webtool where proteins were automatically segmented into 50 aa-sized segments and peptide sequences with their associated intensities were mapped and summed within each. To measure regional differences in peptide yield across protein structures that are independent from changes in whole protein abundance, summed peptide intensities per segment were median normalised based on their corresponding individual protein total summed intensities. Regional age-associated differences in peptide yield across protein structures were visualised by subtracting the average, normalised peptide intensity per segment in the 8-, 52-, and 78-week groups from the 22-week group and dividing by the segment length. Normalised peptide intensities per segment were statistically compared between 8-, 52-, and 78-week groups with the 22-week group using a Bonferroni-corrected, unpaired repeated measure ANOVA. For more information on PLF and its application, please refer to the original MPLF development publication [59].

### Statistical analyses

Data was tested for normality where appropriate using the D’Agostino and Pearson test and statistical tests are indicated in figure legends. Due to non-parametric data, data are presented as median±95% confidence intervals unless otherwise indicated in figure legends. Sample number (n) indicates the number of independent biological samples in each experiment. Sample numbers and experimental repeats are indicated in figures and figure legends or methods section above. Differences in means were considered statistically significant at p<0.05. Significance levels are: \**p*<0.05; \*\**p*<0.01; \*\*\**p*<0.001; \*\*\*\**p*<0.0001. Actual p-values are shown where appropriate. Statistical analyses were performed using the GraphPad Prism 9.5.1 software.

### Disclosure

The authors declare no competing interests.

### Data sharing

The mass spectrometry proteomics data have been deposited to the ProteomeXchange Consortium via the PRIDE partner repository with the dataset identifier PXD070694.

## Acknowledgements

This research was primarily funded by a Kidney Research UK Fellowship to R.P. (TF_007_20181122). R.L. was funded by Kidney Research UK and The Stoneygate Trust (Alport Research Hub), the Wellcome Trust (Ref: 226804/Z/22/Z, Ref: 227417/Z/23/Z, Ref: 301803/Z/23/Z), and the NIHR Manchester Biomedical Research Centre (NIHR203308), which funded E.W. A.H. was funded by a BBSRC CASE DTP studentship and then by the Tissue Biology Platform jointly funded by the Kennedy Trust for Rheumatology Research and Versus Arthritis (KENN 21 22 11). J.S was funded by a BBSRC sLoLa grant (BB/T001984/1). A.E. was funded by a BBSRC New Investigator Award (BB/Z517355/1). J.C. was funded by a UKRI MRC fellowship (MR/W016796/1). The authors would like to thank the University of Manchester Biological Support Facility for assistance in animal welfare. The mass spectrometry-based proteomics was performed by the Biological Mass Spectrometry Core Research Facility in the Faculty of Biology, Medicine and Health (University of Manchester) with the assistance of S Warwood and D Knight (RRID code: SCR_020987).

## References

1. Hynes RO (2009) The extracellular matrix: not just pretty fibrils. Science 326:1216–1219.

2. Lennon R, Byron A, Humphries JD, Randles MJ, Carisey A, Murphy S, Knight D, Brenchley PE, Zent R, Humphries MJ (2014) Global Analysis Reveals the Complexity of the Human Glomerular Extracellular Matrix. Journal of the American Society of Nephrology 25.

3. Ruiz-Ortega M, Rayego-Mateos S, Lamas S, Ortiz A, Rodrigues-Diez RR (2020) Targeting the progression of chronic kidney disease. Nat Rev Nephrol 16:269–288.

4. Glassock RJ, Warnock DG, Delanaye P (2017) The global burden of chronic kidney disease: estimates, variability and pitfalls. Nat Rev Nephrol 13:104–114.

5. Mills KT, Xu Y, Zhang W, Bundy JD, Chen CS, Kelly TN, Chen J, He J (2015) A systematic analysis of worldwide population-based data on the global burden of chronic kidney disease in 2010. Kidney Int 88:950–957.

6. Glassock RJ, Rule AD (2012) The implications of anatomical and functional changes of the aging kidney: with an emphasis on the glomeruli. Kidney international 82:270–277.

7. Kaplan C, Pasternack B, Shah H, Gallo G (1975) Age-related incidence of sclerotic glomeruli in human kidneys. The American journal of pathology 80:227.

8. McLachlan MS (1978) The ageing kidney. Lancet 2:143–145.

9. Alfano G, Perrone R, Fontana F, Ligabue G, Giovanella S, Ferrari A, Gregorini M, Cappelli G, Magistroni R, Donati G (2022) Rethinking Chronic Kidney Disease in the Aging Population. Life 12:1724.

10. Chautard E, Ballut L, Thierry-Mieg N, Ricard-Blum S (2009) MatrixDB, a database focused on extracellular protein–protein and protein–carbohydrate interactions. Bioinformatics 25:690–691.

11. Berthollier C, Vallet SD, Deniaud M, Clerc O, Ricard-Blum S (2021) Building Protein-Protein and Protein-Glycosaminoglycan Interaction Networks Using MatrixDB, the Extracellular Matrix Interaction Database. Current Protocols 1:e47.

12. Naba A, Clauser KR, Hoersch S, Liu H, Carr SA, Hynes RO (2012) The matrisome: in silico definition and in vivo characterization by proteomics of normal and tumor extracellular matrices. Molecular & Cellular Proteomics 11.

13. Shao X, Taha IN, Clauser KR, Gao YT, Naba A (2020) MatrisomeDB: the ECM-protein knowledge database. Nucleic Acids Res 48:D1136–d1144.

14. Jayadev R, Morais M, Ellingford JM, Srinivasan S, Naylor RW, Lawless C, Li AS, Ingham JF, Hastie E, Chi Q, Fresquet M, Koudis NM, Thomas HB, O’Keefe RT, Williams E, Adamson A, Stuart HM, Banka S, Smedley D, Sherwood DR, Lennon R (2022) A basement membrane discovery pipeline uncovers network complexity, regulators, and human disease associations. Sci Adv 8:eabn2265.

15. Naylor RW, Morais M, Lennon R (2021) Complexities of the glomerular basement membrane. Nat Rev Nephrol 17:112–127.

16. Bülow RD, Boor P (2019) Extracellular Matrix in Kidney Fibrosis: More Than Just a Scaffold. J Histochem Cytochem 67:643–661.

17. Michael AF (1984) The glomerular mesangium. Contrib Nephrol 40:7–16.

18. Schaefer L, Schaefer RM (2010) Proteoglycans: from structural compounds to signaling molecules. Cell and tissue research 339:237–246.

19. Border WA, Noble NA, Yamamoto T, Harper JR, Yamaguchi Y, Pierschbacher MD, Ruoslahti E (1992) Natural inhibitor of transforming growth factor-β protects against scarring in experimental kidney disease. Nature 360:361–364.

20. Chen S, Birk DE (2013) The regulatory roles of small leucine-rich proteoglycans in extracellular matrix assembly. The FEBS journal 280:2120–2137.

21. Schaefer L, Iozzo RV (2008) Biological functions of the small leucine-rich proteoglycans: from genetics to signal transduction. Journal of Biological Chemistry 283:21305–21309.

22. Schaefer L, Gröne H-J, Raslik I, Robenek H, Ugorcakova J, Budny S, Schaefer RM, Kresse H (2000) Small proteoglycans of normal adult human kidney: distinct expression patterns of decorin, biglycan, fibromodulin, and lumican. Kidney international 58:1557–1568.

23. Stokes MB, Hudkins KL, Zaharia V, Taneda S, Alpers CE (2001) Up-regulation of extracellular matrix proteoglycans and collagen type I in human crescentic glomerulonephritis. Kidney Int 59:532–542.

24. Stokes MB, Holler S, Cui Y, Hudkins KL, Eitner F, Fogo A, Alpers CE (2000) Expression of decorin, biglycan, and collagen type I in human renal fibrosing disease. Kidney Int 57:487–498.

25. Hägg PM, Hägg PO, Peltonen S, Autio-Harmainen H, Pihlajaniemi T (1997) Location of type XV collagen in human tissues and its accumulation in the interstitial matrix of the fibrotic kidney. Am J Pathol 150:2075–2086.

26. Zaferani A, Talsma DT, Yazdani S, Celie JW, Aikio M, Heljasvaara R, Navis GJ, Pihlajaniemi T, van den Born J (2014) Basement membrane zone collagens XV and XVIII/proteoglycans mediate leukocyte influx in renal ischemia/reperfusion. PLoS One 9:e106732.

27. Bretaud S, Guillon E, Karppinen SM, Pihlajaniemi T, Ruggiero F (2020) Collagen XV, a multifaceted multiplexin present across tissues and species. Matrix Biol Plus 6-7:100023.

28. Clotet-Freixas S, Konvalinka A (2021) Too Little or Too Much? Extracellular Matrix Remodeling in Kidney Health and Disease. J Am Soc Nephrol 32:1541–1543.

29. Bonnans C, Chou J, Werb Z (2014) Remodelling the extracellular matrix in development and disease. Nature Reviews Molecular Cell Biology 15:786–801.

30. Eckersley A, Yamamura T, Lennon R (2023) Matrikines in kidney ageing and age-related disease. Current Opinion in Nephrology and Hypertension 32:551–558.

31. Randles MJ, Humphries MJ, Lennon R (2017) Proteomic definitions of basement membrane composition in health and disease. Matrix Biol 57-58:12–28.

32. Louzao-Martinez L, Van Dijk CG, Xu YJ, Korn A, Bekker NJ, Brouwhuis R, Nicese MN, Demmers JA, Goumans M-JT, Masereeuw R (2019) A proteome comparison between human fetal and mature renal extracellular matrix identifies EMILIN1 as a regulator of renal epithelial cell adhesion. Matrix biology plus 4:100011.

33. Hobeika L, Barati MT, Caster DJ, McLeish KR, Merchant ML (2017) Characterization of glomerular extracellular matrix by proteomic analysis of laser-captured microdissected glomeruli. Kidney international 91:501–511.

34. Rende U, Ahn SB, Adhikari S, Moh ESX, Pollock CA, Saad S, Guller A (2023) Deciphering the Kidney Matrisome: Identification and Quantification of Renal Extracellular Matrix Proteins in Healthy Mice. International Journal of Molecular Sciences 24:2827.

35. Randles MJ, Woolf AS, Huang JL, Byron A, Humphries JD, Price KL, Kolatsi-Joannou M, Collinson S, Denny T, Knight D (2015) Genetic background is a key determinant of glomerular extracellular matrix composition and organization. Journal of the American Society of Nephrology: JASN 26:3021.

36. Paunas FTI, Finne K, Leh S, Osman TA-H, Marti H-P, Berven F, Vikse BE (2019) Characterization of glomerular extracellular matrix in IgA nephropathy by proteomic analysis of laser-captured microdissected glomeruli. BMC nephrology 20:1–12.

37. Merchant ML, Barati MT, Caster DJ, Hata JL, Hobeika L, Coventry S, Brier ME, Wilkey DW, Li M, Rood IM, Deegens JK, Wetzels JF, Larsen CP, Troost JP, Hodgin JB, Mariani LH, Kretzler M, Klein JB, McLeish KR (2020) Proteomic Analysis Identifies Distinct Glomerular Extracellular Matrix in Collapsing Focal Segmental Glomerulosclerosis. J Am Soc Nephrol 31:1883–1904.

38. Randles MJ, Lausecker F, Kong Q, Suleiman H, Reid G, Kolatsi-Joannou M, Davenport B, Tian P, Falcone S, Potter P, Van Agtmael T, Norman JT, Long DA, Humphries MJ, Miner JH, Lennon R (2021) Identification of an Altered Matrix Signature in Kidney Aging and Disease. J Am Soc Nephrol 32:1713–1732.

39. Lipp SN, Jacobson KR, Hains DS, Schwarderer AL, Calve S (2021) 3D Mapping Reveals a Complex and Transient Interstitial Matrix During Murine Kidney Development. Journal of the American Society of Nephrology 32:1649–1665.

40. Lausecker F, Lennon R, Randles MJ (2022) The kidney matrisome in health, aging, and disease. Kidney International 102:1000–1012.

41. Vilchez D, Saez I, Dillin A (2014) The role of protein clearance mechanisms in organismal ageing and age-related diseases. Nat Commun 5:5659.

42. Labbadia J, Morimoto RI (2015) The biology of proteostasis in aging and disease. Annu Rev Biochem 84:435–464.

43. Kennedy BK, Berger SL, Brunet A, Campisi J, Cuervo AM, Epel ES, Franceschi C, Lithgow GJ, Morimoto RI, Pessin JE, Rando TA, Richardson A, Schadt EE, Wyss-Coray T, Sierra F (2014) Geroscience: linking aging to chronic disease. Cell 159:709–713.

44. López-Otín C, Blasco MA, Partridge L, Serrano M, Kroemer G (2013) The hallmarks of aging. Cell 153:1194–1217.

45. Selman M, Pardo A (2021) Fibroageing: An ageing pathological feature driven by dysregulated extracellular matrix-cell mechanobiology. Ageing Research Reviews 70:101393.

46. Decaris ML, Gatmaitan M, FlorCruz S, Luo F, Li K, Holmes WE, Hellerstein MK, Turner SM, Emson CL (2014) Proteomic analysis of altered extracellular matrix turnover in bleomycin-induced pulmonary fibrosis. Mol Cell Proteomics 13:1741–1752.

47. Trier JS, Allan CH, Abrahamson DR, Hagen SJ (1990) Epithelial basement membrane of mouse jejunum. Evidence for laminin turnover along the entire crypt-villus axis. J Clin Invest 86:87–95.

48. Matsubayashi Y, Sanchez-Sanchez BJ, Marcotti S, Serna-Morales E, Dragu A, Diaz-de-la-Loza MD, Vizcay-Barrena G, Fleck RA, Stramer BM (2020) Rapid Homeostatic Turnover of Embryonic ECM during Tissue Morphogenesis. Dev Cell 54:33–42 e39.

49. Keeley DP, Hastie E, Jayadev R, Kelley LC, Chi Q, Payne SG, Jeger JL, Hoffman BD, Sherwood DR (2020) Comprehensive Endogenous Tagging of Basement Membrane Components Reveals Dynamic Movement within the Matrix Scaffolding. Dev Cell 54:60–74 e67.

50. Mao M, Ishikawa Y, Labelle-Dumais C, Wang X, Kuo YM, Gaffney UB, Smith ME, Abdala CN, Lebedev MD, Paradee WJ, Gould DB (2025) A multifunction murine Col4a1 allele reveals potential gene therapy parameters for Gould syndrome. J Cell Biol 224.

51. Liu P, Xie X, Jin J (2020) Isotopic Nitrogen-15 Labeling of Mice Identified Long-lived Proteins of the Renal Basement Membranes. Sci Rep 10:5317.

52. Hinkson IV, Elias JE (2011) The dynamic state of protein turnover: It’s about time. Trends Cell Biol 21:293–303.

53. Rolfs Z, Frey BL, Shi X, Kawai Y, Smith LM, Welham NV (2021) An atlas of protein turnover rates in mouse tissues. Nat Commun 12:6778.

54. Krüger M, Moser M, Ussar S, Thievessen I, Luber CA, Forner F, Schmidt S, Zanivan S, Fässler R, Mann M (2008) SILAC mouse for quantitative proteomics uncovers kindlin-3 as an essential factor for red blood cell function. Cell 134:353–364.

55. Ng SS, Park JE, Meng W, Chen CP, Kalaria RN, McCarthy NE, Sze SK (2020) Pulsed SILAM Reveals In Vivo Dynamics of Murine Brain Protein Translation. ACS Omega 5:13528–13540.

56. Fornasiero EF, Mandad S, Wildhagen H, Alevra M, Rammner B, Keihani S, Opazo F, Urban I, Ischebeck T, Sakib MS, Fard MK, Kirli K, Centeno TP, Vidal RO, Rahman R-U, Benito E, Fischer A, Dennerlein S, Rehling P, Feussner I, Bonn S, Simons M, Urlaub H, Rizzoli SO (2018) Precisely measured protein lifetimes in the mouse brain reveal differences across tissues and subcellular fractions. Nature Communications 9:4230.

57. Ariosa-Morejon Y, Santos A, Fischer R, Davis S, Charles P, Thakker R, Wann AK, Vincent TL (2021) Age-dependent changes in protein incorporation into collagen-rich tissues of mice by in vivo pulsed SILAC labelling. Elife 10.

58. Eckersley A, Ozols M, O’Cualain R, Keevill E-J, Foster A, Pilkington S, Knight D, Griffiths CE, Watson RE, Sherratt MJ (2020) Proteomic fingerprints of damage in extracellular matrix assemblies. Matrix Biology Plus 5:100027.

59. Ozols M, Eckersley A, Mellody KT, Mallikarjun V, Warwood S, O’Cualain R, Knight D, Watson RE, Griffiths CE, Swift J (2021) Peptide location fingerprinting reveals modification-associated biomarker candidates of ageing in human tissue proteomes. Aging Cell 20:e13355.

60. Eckersley A, Ozols M, Chen P, Tam V, Hoyland JA, Trafford A, Chan D, Sherratt MJ (2021) Peptide location fingerprinting reveals tissue region-specific differences in protein structures in an ageing human organ. International Journal of Molecular Sciences 22:10408.

61. Eckersley A, Ozols M, Chen P, Tam V, Ward LJ, Hoyland JA, Trafford A, Yuan X-M, Schiller HB, Chan D (2022) Peptide location fingerprinting identifies species-and tissue-conserved structural remodelling of proteins as a consequence of ageing and disease. Matrix Biology 114:108–137.

62. Brand O, Kirkham S, Jagger C, Ozols M, Purohit K, Zhang Z, Lennon R, Hussell T, Eckersley A (2025) Lung basement membranes are compositionally and structurally altered following resolution of influenza infection. Mucosal Immunology.

63. Eckersley A, Morais MRPT, Ozols M, Lennon R (2023) Peptide location fingerprinting identifies structural alterations within basement membrane components in ageing kidney. Matrix Biology 121:167–178.

64. Baghy K, Szakadáti H, Kovalszky I (2025) Decorin the antifibrotic proteoglycan and its progression in therapy. American Journal of Physiology-Cell Physiology 328:C1853–C1865.

65. Genovese F, Manresa AA, Leeming DJ, Karsdal MA, Boor P (2014) The extracellular matrix in the kidney: a source of novel non-invasive biomarkers of kidney fibrosis? Fibrogenesis Tissue Repair 7:4.

66. Alevra M, Mandad S, Ischebeck T, Urlaub H, Rizzoli SO, Fornasiero EF (2019) A mass spectrometry workflow for measuring protein turnover rates in vivo. Nat Protoc 14:3333–3365.

67. Johnson TS, Fisher M, Haylor JL, Hau Z, Skill NJ, Jones R, Saint R, Coutts I, Vickers ME, El Nahas AM, Griffin M (2007) Transglutaminase Inhibition Reduces Fibrosis and Preserves Function in Experimental Chronic Kidney Disease. Journal of the American Society of Nephrology 18:3078–3088.

68. Arnold P, Otte A, Becker-Pauly C (2017) Meprin metalloproteases: Molecular regulation and function in inflammation and fibrosis. Biochimica et Biophysica Acta (BBA) - Molecular Cell Research 1864:2096–2104.

69. Prox J, Arnold P, Becker-Pauly C (2015) Meprin α and meprin β: Procollagen proteinases in health and disease. Matrix Biology 44-46:7–13.

70. Kruppa D, Peters F, Bornert O, Maler MD, Martin SF, Becker-Pauly C, Nyström A (2021) Distinct contributions of meprins to skin regeneration after injury - Meprin α a physiological processer of pro-collagen VII. Matrix Biol Plus 11:100065.

71. Herzog C, Haun RS, Kaushal GP (2019) Role of meprin metalloproteinases in cytokine processing and inflammation. Cytokine 114:18–25.

72. Bayly-Jones C, Lupton CJ, Fritz C, Venugopal H, Ramsbeck D, Wermann M, Jäger C, de Marco A, Schilling S, Schlenzig D, Whisstock JC (2022) Helical ultrastructure of the metalloprotease meprin α in complex with a small molecule inhibitor. Nature Communications 13:6178.

73. Ishmael FT, Norcum MT, Benkovic SJ, Bond JS (2001) Multimeric structure of the secreted meprin A metalloproteinase and characterization of the functional protomer. J Biol Chem 276:23207–23211.

74. Sterchi EE, Stöcker W, Bond JS (2008) Meprins, membrane-bound and secreted astacin metalloproteinases. Mol Aspects Med 29:309–328.

75. Ishmael SS, Ishmael FT, Jones AD, Bond JS (2006) Protease domain glycans affect oligomerization, disulfide bond formation, and stability of the meprin A metalloprotease homo-oligomer. J Biol Chem 281:37404–37415.

76. Arolas JL, Broder C, Jefferson T, Guevara T, Sterchi EE, Bode W, Stöcker W, Becker-Pauly C, Gomis-Rüth FX (2012) Structural basis for the sheddase function of human meprin β metalloproteinase at the plasma membrane. Proceedings of the National Academy of Sciences 109:16131–16136.

77. Peters F, Scharfenberg F, Colmorgen C, Armbrust F, Wichert R, Arnold P, Potempa B, Potempa J, Pietrzik CU, Häsler R, Rosenstiel P, Becker-Pauly C (2019) Tethering soluble meprin α in an enzyme complex to the cell surface affects IBD-associated genes. Faseb j 33:7490–7504.

78. Jefferson T, Auf dem Keller U, Bellac C, Metz VV, Broder C, Hedrich J, Ohler A, Maier W, Magdolen V, Sterchi E (2013) The substrate degradome of meprin metalloproteases reveals an unexpected proteolytic link between meprin β and ADAM10. Cellular and Molecular Life Sciences 70:309–333.

79. Herzog C, Haun RS, Ludwig A, Shah SV, Kaushal GP (2014) ADAM10 is the major sheddase responsible for the release of membrane-associated meprin A. Journal of Biological Chemistry 289:13308–13322.

80. Miner JH (2012) The glomerular basement membrane. Exp Cell Res 318:973–978.

81. Aumailley M, Battaglia C, Mayer U, Reinhardt D, Nischt R, Timpl R, Fox JW (1993) Nidogen mediates the formation of ternary complexes of basement membrane components. Kidney International 43:7–12.

82. Töpfer U, Holz A (2024) Nidogen in development and disease. Frontiers in Cell and Developmental Biology Volume 12–2024.

83. Dai J, Estrada B, Jacobs S, Sánchez-Sánchez BJ, Tang J, Ma M, Magadán-Corpas P, Pastor-Pareja JC, Martín-Bermudo MD (2018) Dissection of Nidogen function in Drosophila reveals tissue-specific mechanisms of basement membrane assembly. PLoS Genetics 14:e1007483.

84. Reinhardt D, Mann K, Nischt R, Fox J, Chu M, Krieg T, Timpl R (1993) Mapping of nidogen binding sites for collagen type IV, heparan sulfate proteoglycan, and zinc. Journal of Biological Chemistry 268:10881–10887.

85. Hopf M, Göhring W, Mann K, Timpl R (2001) Mapping of binding sites for nidogens, fibulin-2, fibronectin and heparin to different IG modules of perlecan. Journal of molecular biology 311:529–541.

86. Gerl M, Mann K, Aumailley M, Timpl R (1991) Localization of a major nidogen-binding site to domain III of laminin B2 chain. European journal of biochemistry 202:167–174.

87. Godwin ARF, Becker MH, Dajani R, Snee M, Roseman AM, Baldock C (2025) Collagen VI microfibril structure reveals mechanism for molecular assembly and clustering of inherited pathogenic mutations. Nat Commun 16:7549.

88. Bock F, Li S, Pozzi A, Zent R (2025) Integrins in the kidney — beyond the matrix. Nature Reviews Nephrology 21:157–174.

89. Pfaff M, Aumailley M, Specks U, Knolle J, Zerwes HG, Timpl R (1993) Integrin and Arg-Gly-Asp dependence of cell adhesion to the native and unfolded triple helix of collagen type VI. Exp Cell Res 206:167–176.

90. Wirz JA, Boudko SP, Lerch TF, Chapman MS, Bächinger HP (2011) Crystal structure of the human collagen XV trimerization domain: a potent trimerizing unit common to multiplexin collagens. Matrix Biol 30:9–15.

91. Vleming L, Baelde J, Westendorp R, Daha M, Van Es L, Bruijn J (1995) Progression of chronic renal disease in humans is associated with the deposition of basement membrane components and decorin in the interstitial extracellular matrix. Clinical nephrology 44:211–219.

92. Mason RM, Wahab NA (2003) Extracellular matrix metabolism in diabetic nephropathy. Journal of the American Society of Nephrology 14:1358–1373.

93. Rasmussen DGK, Fenton A, Jesky M, Ferro C, Boor P, Tepel M, Karsdal MA, Genovese F, Cockwell P (2017) Urinary endotrophin predicts disease progression in patients with chronic kidney disease. Scientific reports 7:17328.

94. Sparding N, Genovese F, Rasmussen DGK, Karsdal MA, Neprasova M, Maixnerova D, Satrapova V, Frausova D, Hornum M, Bartonova L (2022) Endotrophin, a collagen type VI-derived matrikine, reflects the degree of renal fibrosis in patients with IgA nephropathy and in patients with ANCA-associated vasculitis. Nephrology Dialysis Transplantation 37:1099–1108.

95. Fenton A, Jesky MD, Ferro CJ, Sørensen J, Karsdal MA, Cockwell P, Genovese F (2017) Serum endotrophin, a type VI collagen cleavage product, is associated with increased mortality in chronic kidney disease. PLoS One 12:e0175200.

96. Lin CHS, Chen J, Ziman B, Marshall S, Maizel J, Goligorsky MS (2014) Endostatin and kidney fibrosis in aging: a case for antagonistic pleiotropy? American Journal of Physiology-Heart and Circulatory Physiology 306:H1692–H1699.

97. Lin CHS, Chen J, Zhang Z, Johnson GV, Cooper AJ, Feola J, Bank A, Shein J, Ruotsalainen HJ, Pihlajaniemi TA (2016) Endostatin and transglutaminase 2 are involved in fibrosis of the aging kidney. Kidney international 89:1281–1292.

98. Soylemezoglu O, Wild G, Dalley AJ, MacNeil S, Milford-Ward A, Brown CB, el Nahas AM (1997) Urinary and serum type III collagen: markers of renal fibrosis. Nephrol Dial Transplant 12:1883–1889.

99. Mavrogeorgis E, Mischak H, Latosinska A, Vlahou A, Schanstra JP, Siwy J, Jankowski V, Beige J, Jankowski J (2021) Collagen-Derived Peptides in CKD: A Link to Fibrosis. Toxins (Basel) 14.

100. Tatsukawa H, Otsu R, Tani Y, Wakita R, Hitomi K (2018) Isozyme-specific comprehensive characterization of transglutaminase-crosslinked substrates in kidney fibrosis. Sci Rep 8:7306.

101. Schaefer L (2011) Small leucine-rich proteoglycans in kidney disease. J Am Soc Nephrol 22:1200–1207.

102. Meng X-m, Nikolic-Paterson DJ, Lan HY (2016) TGF-β: the master regulator of fibrosis. Nature Reviews Nephrology 12:325–338.

103. Zhou S, Yin X, Mayr M, Noor M, Hylands PJ, Xu Q (2020) Proteomic landscape of TGF-β1-induced fibrogenesis in renal fibroblasts. Scientific Reports 10:19054.

104. Schnaper HW, Jandeska S, Runyan CE, Hubchak SC, Basu RK, Curley JF, Smith RD, Hayashida T (2009) TGF-beta signal transduction in chronic kidney disease. Front Biosci (Landmark Ed) 14:2448–2465.

105. Broder C, Arnold P, Vadon-Le Goff S, Konerding MA, Bahr K, Müller S, Overall CM, Bond JS, Koudelka T, Tholey A (2013) Metalloproteases meprin α and meprin β are C-and N-procollagen proteinases important for collagen assembly and tensile strength. Proceedings of the National Academy of Sciences 110:14219–14224.

106. Kronenberg D, Bruns BC, Moali C, Vadon-Le Goff S, Sterchi EE, Traupe H, Böhm M, Hulmes DJ, Stöcker W, Becker-Pauly C (2010) Processing of procollagen III by meprins: new players in extracellular matrix assembly? Journal of investigative dermatology 130:2727–2735.

107. Herzog C, Marisiddaiah R, Haun RS, Kaushal GP (2015) Basement membrane protein nidogen-1 is a target of meprin β in cisplatin nephrotoxicity. Toxicol Lett 236:110–116.

108. Bertenshaw GP, Norcum MT, Bond JS (2003) Structure of homo- and hetero-oligomeric meprin metalloproteases. Dimers, tetramers, and high molecular mass multimers. J Biol Chem 278:2522–2532.

