## Supplementary Materials for "Ageing impacts extracellular matrix turnover and remodelling in the kidney"

***Corresponding author email address***

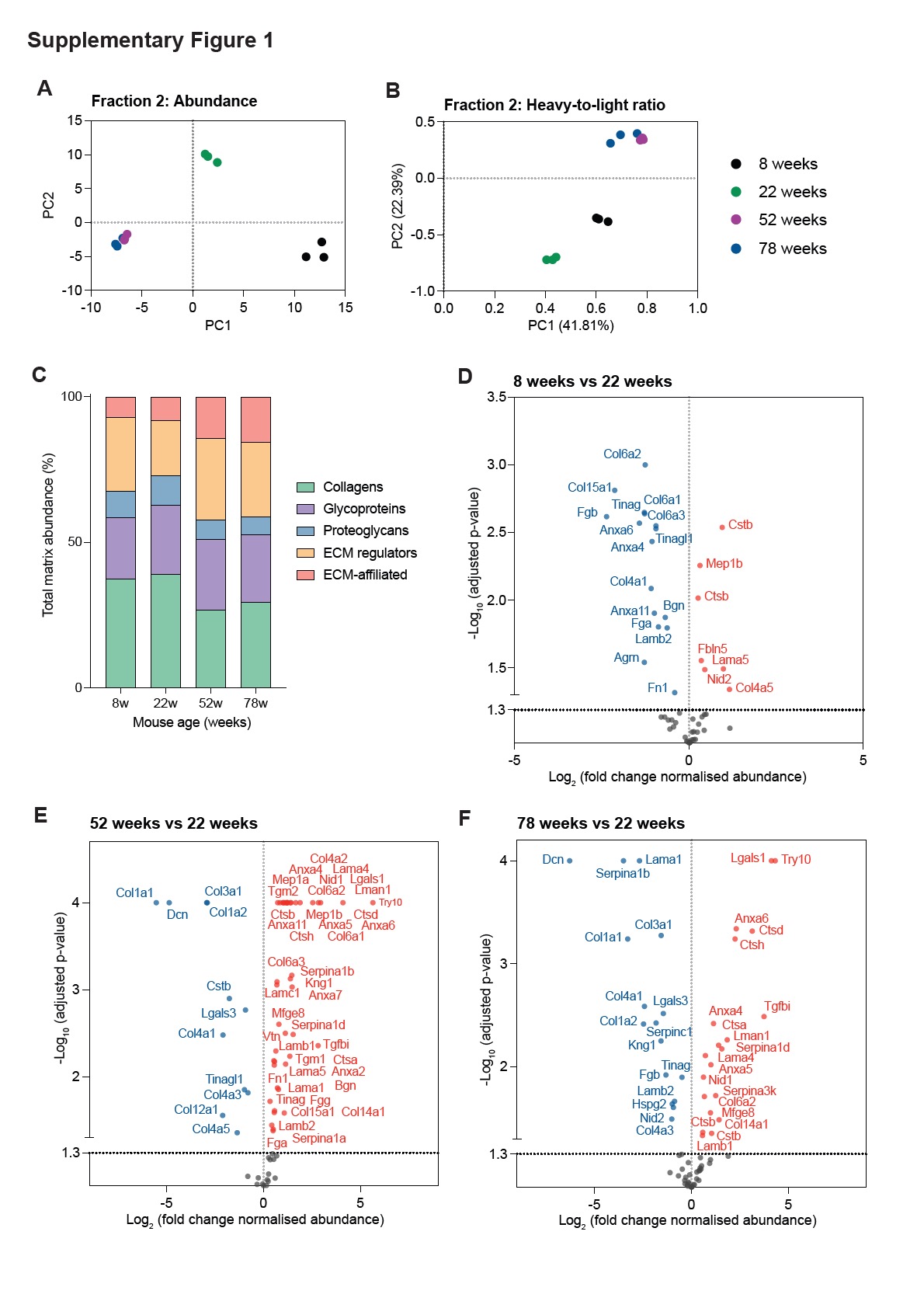

**Supplementary Figure 1:** Principal components analysis (PCA) of **A**) normalised abundance, and **B**) heavy-to-light ratios from the kidney matrix-enriched fraction from 8-, 22-, 52-, and 78-week mice. **C**) Distribution of total kidney matrix abundance derived from heavy and light MS1 ion intensities from matrisome proteins in the matrix-enriched fraction from 8-, 22-, 52- and 78-week-old mice demonstrating a relative reduction in core matrix proteins with age. **D**) Volcano plot showing fold-changes in abundance of kidney matrix proteins identified in the matrix-enriched fraction from 8- week compared to 22-week mice. **E**) Volcano plot showing fold-changes in abundance of kidney matrix proteins identified in the matrix-enriched fraction from 52- week compared to 22-week mice. **F**) Volcano plot showing fold-changes in abundance of kidney matrix proteins identified in the matrix-enriched fraction from 78- week compared to 22-week mice. For (D), (E), and (F), matrix proteins with significantly increased abundance in 8-, 52-, or 78-week mice are shown in red and matrix proteins with significantly decreased abundance in 8-, 52-, or 78-week mice are shown in blue. Dotted black line represents significance threshold (adjusted p<0.05). N=3 biological replicates per condition for all analysis.

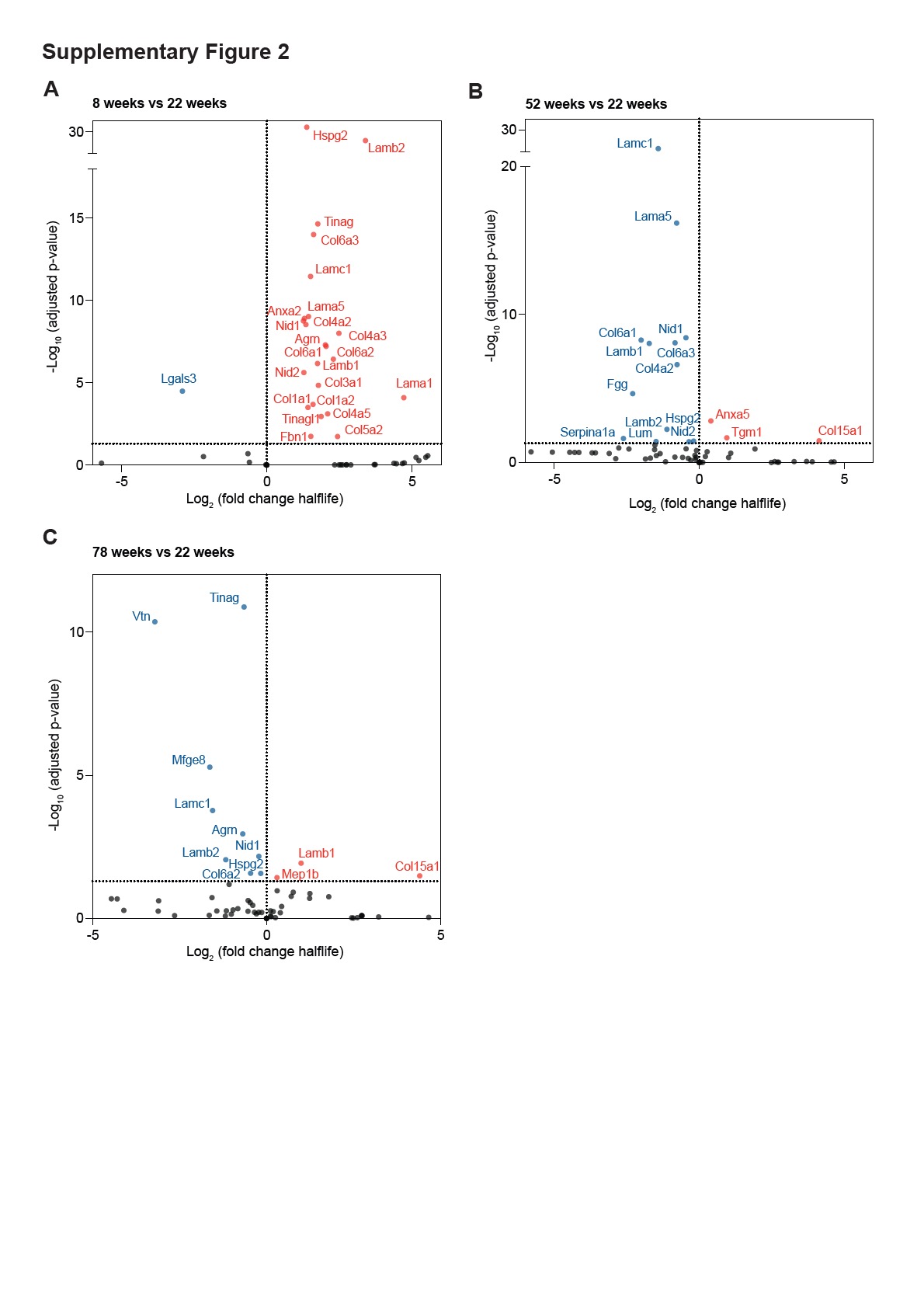

**Supplementary Figure 2:** Volcano plot showing fold changes in half-lives of kidney matrix proteins identified in the matrix-enriched fraction from **A**) 8-week vs 22-week, **B**) 52-week vs 22-week, and **C**) 78-week vs 22-week mice. Matrix proteins with significantly shorter half-lives (faster turnover) in 8-, 52-, or 78-week mice are shown in red and matrix proteins with significantly longer half-lives (slower turnover) in 8-, 52-, or 78-week mice are shown in blue. Dotted black line represents significance threshold (adjusted p<0.05). N=3 biological replicates per condition for all analysis.

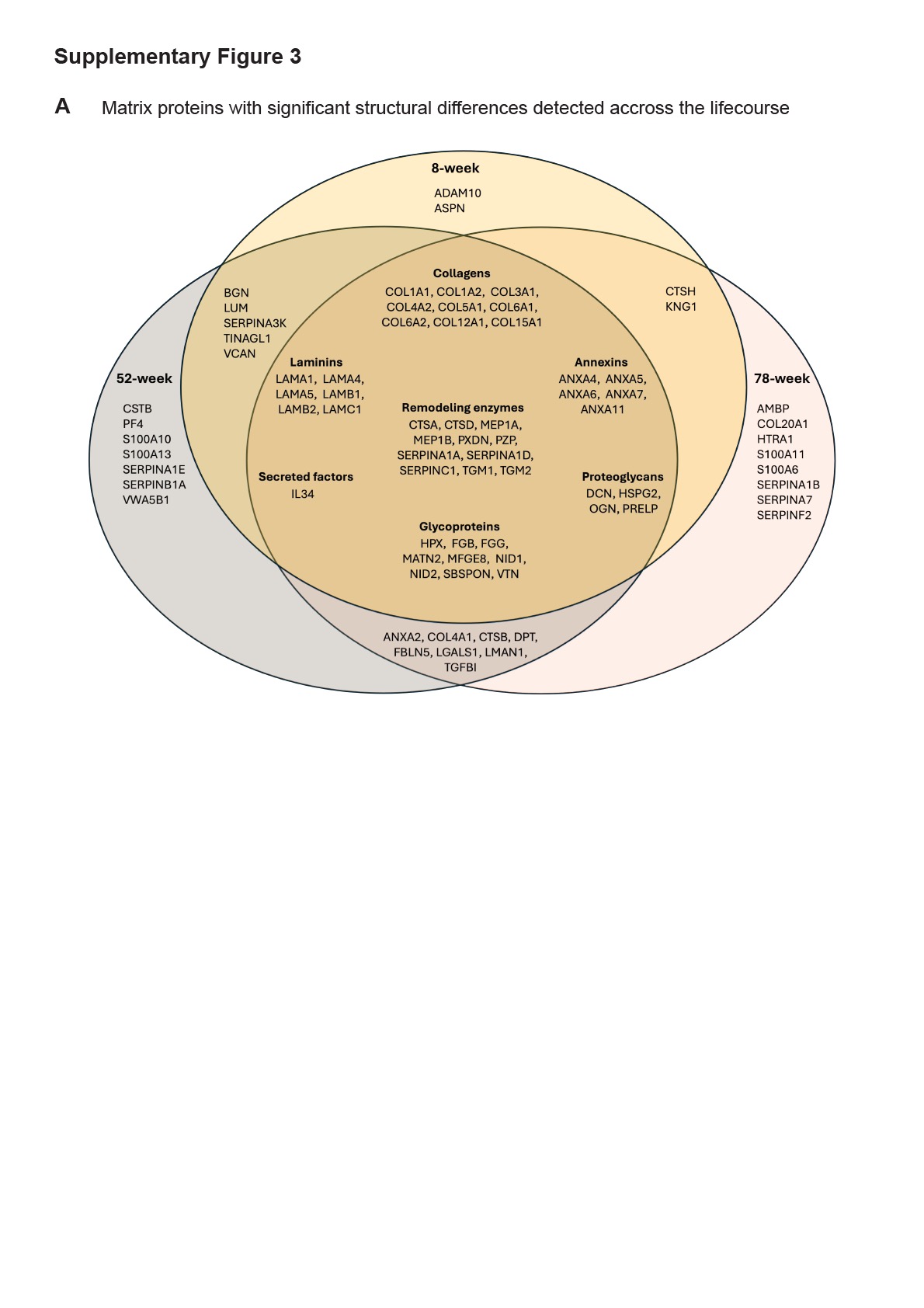

**Supplementary Figure 3: A**) Kidney matrix proteins identified by PLF analysis with statistically significantly structure-associated differences between 8-week (middle), 52-week (left) and 78-week (right) compared to 22-week. Central overlap shows a total of 45 matrix proteins were identified as significantly structurally different in 8-, 52-, and 78-week mice relative to 22-week mice.
